# Profiles of expressed mutations in single cells reveal subclonal expansion patterns and therapeutic impact of intratumor heterogeneity

**DOI:** 10.1101/2021.03.26.437185

**Authors:** Farid Rashidi Mehrabadi, Kerrie L. Marie, Eva Pérez-Guijarro, Salem Malikić, Erfan Sadeqi Azer, Howard H. Yang, Can Kızılkale, Charli Gruen, Welles Robinson, Huaitian Liu, Michael C. Kelly, Christina Marcelus, Sandra Burkett, Aydın Buluç, Funda Ergün, Maxwell P. Lee, Glenn Merlino, Chi-Ping Day, S. Cenk Sahinalp

## Abstract

Advances in single-cell RNA sequencing (scRNAseq) technologies uncovered an unexpected complexity in tumors, underlining the relevance of intratumor heterogeneity to cancer progression and therapeutic resistance. Heterogeneity in the mutational composition of cancer cells is a result of distinct (sub)clonal expansions, each with a distinct metastatic potential and resistance to specific treatments. Unfortunately, due to their low read coverage per cell, scRNAseq datasets are too sparse and noisy to be used for detecting expressed mutations in single cells. Additionally, the large number of cells and mutations present in typical scRNAseq datasets are too large for available computational tools to, e.g., infer distinct subclones, lineages or trajectories in a tumor. Finally, there are no principled methods to assess distinct subclones inferred through single-cell sequencing data and the genomic alterations that seed and potentially cause them. Here we present `Trisicell`, a computational toolkit for scalable mutational intratumor heterogeneity inference and assessment from scRNAseq as well as single-cell genome or exome sequencing data. `Trisicell` allows reliable identification of distinct clonal lineages of a tumor, offering the ability to focus on the most important subclones and the genomic alterations that are associated with tumor proliferation. We comprehensively assessed `Trisicell` on a melanoma model by comparing distinct lineages and subclones it identifies on scRNAseq data, to those inferred using matching bulk whole exome (bWES) and transcriptome (bWTS) sequencing data from clonal sublines derived from single cells. Our results demonstrate that distinct lineages and subclones of a tumor can be reliably inferred and evaluated based on mutation calls from scRNAseq data through the use of `Trisicell`. Additionally, they reveal a strong correlation between aggressiveness and mutational composition, both across the inferred subclones, and among human melanomas. We also applied `Trisicell` to infer and evaluate distinct subclonal expansion patterns of the same mouse melanoma model after treatment with immune checkpoint blockade (ICB). After integratively analyzing our cell-specific mutation calls with their expression profiles, we observed that each subclone with a distinct set of novel somatic mutations is strongly associated with a specific developmental status. Moreover, each subclone had developed a unique ICB-resistance mechanism. These results demonstrate that `Trisicell` can robustly utilize scRNAseq data to delineate intratumor heterogeneity and help understand biological mechanisms underlying tumor progression and resistance to therapy.

## 1 Introduction

Cancer is a genomic disease that originates in distinct cells and grows through a complex process involving acquisition of genomic alterations, environmental selection, clonal expansion, and competition. The “natural history” of cancer development results in extensive intratumor heterogeneity, which drives disease progression and therapeutic resistance [1, 2, 3, 4]. From a genomic point of view, the subclonal makeup of tumors can be better understood through modeling their progression as a branched process, in the form of a clonal or mutation tree^1^ [6, 7, 8]. These trees depict relative temporal order of genomic alteration events during tumorigenesis and, as such, they can help identify key mutations driving the formation of distinct subpopulations, especially those that are resistant to therapies or have metastatic seeding potential [9].

The first computational tools for reconstructing the subclonal composition of a tumor were developed for bulk genomic sequencing data (see [10] for a survey). Unfortunately, since bulk sequencing provides only an aggregate signal over a large number of cells, it offers a very limited resolution in the inferred clonal tree. Specifically, computational methods based on bulk sequencing data are frequently unable to distinguish between multiple distinct trees that describe the observed data equally well [11, 12, 13].

The rise of single-cell sequencing (SCS) has made it possible to explore the makeup of tumors at a higher, cellular resolution. In particular, single-cell transcriptomic sequencing (scRNAseq) has significantly improved our understanding of intratumor heterogeneity from a gene expression point of view (see [14] for a survey). As per its application to developmental biology, scRNAseq has enabled the identification of tumor subpopulations based on their gene expression, differentiation status and lineages, and how they respond to therapies. Unfortunately, computational methods for tracing lineages through gene expression profiles require a hypothesis of developmental origin, and as such, can not help explore mutational intratumor heterogeneity [15]. In contrast, recently developed barcoding techniques (see [16] for a survey) can help perform detailed lineage tracing up to the time of the original tumor biopsy, but not earlier. Finally, mutation trees inferred based on genomic alterations observed in single cells provide a very detailed progression history of a tumor and are our main focus in this paper.

Many of the methods developed for studying the progression history of a tumor from single-cell data primarily employed genomic mutations and short indels observed in single cells. The first methods in this domain, such as SCITE [7] and OncoNEM [17], aimed to build the “most likely” mutation/clonal tree through genomic SCS data under the “infinite sites” assumption (ISA). Even though some (e.g. SiFit [8] and SiCloneFit [18]) relaxed this assumption by allowing ISA violations at a given “cost” and others (e.g. SPhyR [19]) generalized it to a Dollo parsimony, the majority of available tools still employ ISA (see [20] for a survey), and aim to infer mutation/clonal tree by “correcting for noise” in genotype calls (due to, e.g., allele dropout or low sequence coverage), possibly with a maximum likelihood approach. Since this is a computationally hard (in fact “NP-hard”) problem [21, 22, 23, 19] any principled approach has a theoretical upper bound with respect to the scale of the SCS data it can handle. As a result, available tools either employ MCMC (Markov Chain Monte Carlo) methods and other heuristics (e.g. SCITE [7]), which can not guarantee optimality, or use Integer Linear Programming or Boolean Constraint Satisfaction problem solvers (e.g. PhISCS [24]), which have worst-case running times exponential with the input size. Even recent, custom-built branch and bound approaches such as PhISCS-BnB [25] can not cope with the scale of emerging SCS datasets with thousands of mutations detected in hundreds of cells. As a result, the only scalable methods in existence for this purpose are the “distance-based” heuristics (e.g. neighbor-join-based ScisTree [23]), which have known theoretical limitations and can not offer performance guarantees.^2^ An orthogonal set of tools utilize single-cell copy number variation (CNV) calls, (e.g. offered by Chromium Single Cell CNV Solution from 10x Genomics), to infer the progression history of tumors (e.g. [26]) but they work only with a limited number of (sub)clones and copy number alteration events.

The combination of single-cell sequencing and tumor progression analysis is the key to understanding intratumor heterogeneity. Unfortunately, single-cell whole genome sequencing (scWGS) and single-cell whole exome sequencing (scWES) are prohibitively expensive and suffer from high levels of sequencing noise [27]; as a result, they are not commonly employed. In contrast, single-cell whole transcriptome sequencing protocols, such as Smart-seq2 [28], have gained popularity due to their relatively modest cost. However, their ability to offer information about (expressed) mutations and copy number alterations has been underutilized. In fact, there have been only two attempts to (1) cluster single cells based on their mutational profiles derived from scRNAseq data and/or (2) utilize these clusters to build mutation/clonal trees. The first was DENDRO [29], which, as per ScisTree, is a greedy hierarchical clustering method based on coverage-weighted differences between cells, and thus has theoretical limitations with respect to accuracy. Another was Cardelino [30], which, as per SCITE, simultaneously assigns cells and mutations to nodes/clones. Since Cardelino is based on Gibbs sampling, it offers no performance guarantee and does not scale up to infer sizable/detailed trees (e.g., most of the trees reported in [30] consist of 4 nodes).

To address some of the key challenges in reconstructing the progression history of a tumor from single-cell genomic and, in particular, transcriptomic sequencing data, we developed `Trisicell` (<u>Tri</u>ple-toolkit for <u>si</u>ngle-<u>cell</u> intratumor heterogeneity inference), comprised of three computational methods of independent but complementary aims and applications (see Figure 1): (1) `Trisicell-PartF` is the first method devised to compute the *partition function* of a given mutation tree. Given an input genotype matrix, `Trisicell-PartF` employs a localized sampling strategy to compute the probability of (and thus our confidence in) any user-specified set of cells forming a subtree/clade seeded by one or more (again user-specified) mutations. Importantly, we prove that, as the sample size increases, the output of `Trisicell-PartF` converges to the correct value of the partition function. (2) `Trisicell-Cons` is devised to compare two or more mutation trees derived from the same single-cell data, and build their consensus tree by “collapsing” the minimum number of their edges so that the resulting trees are isomorphic. `Trisicell-Cons` guarantees to build the optimal consensus tree across multiple trees inferred by distinct methods and/or data types, and helps in identifying robust ancestor/descendant relationships between cells and mutations. (3) `Trisicell-Boost` is a booster for tree inference methods allowing them to scale up and handle large and noisy scRNAseq datasets. For that, `Trisicell-Boost`, iteratively samples random subsets of mutations and builds the mutation tree on each subset through a user-selected inference method (e.g. SCITE or PhISCS). It then establishes the consensus of these trees as the global tree that includes the entire set of mutations detected in the tumor sample. In combination, the three components of `Trisicell` provide means to infer large scale trees of tumor progression, compare them in a principled manner, and assess their distinct subtrees/clades, especially through the use of scRNAseq data.

**Figure 1.**
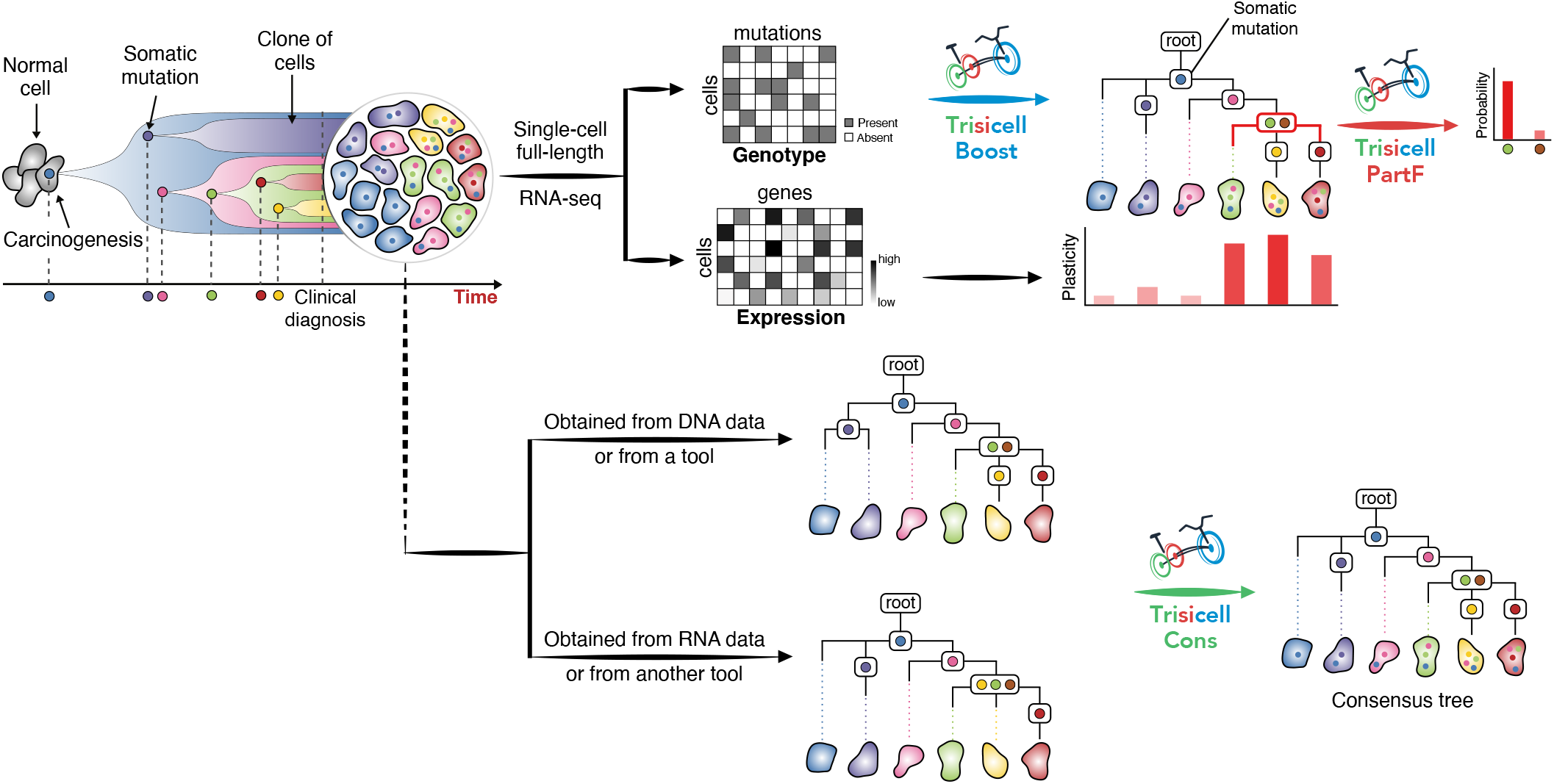
Overview of Trisicell. Tumors are typically heterogeneous and are made of distinct populations of cells, each with a distinct set of somatic mutations. Whole single-cell RNAseq (scRNAseq) data from a tumor is used for calling expressed mutations in each cell; scRNAseq data can be additionally used to derive single-cell expression profiles. `Trisicell-Boost` can then be applied to cell-specific mutation sets for scalable and robust reconstruction of the history of tumor progression. Additional tools may report alternative trees of tumor progression, which can all be integrated into a single consensus tree through the use of `Trisicell-Cons`.^*a*^ Expression profiles of cells harbored in distinct subtrees of the resulting tree may suggest distinct subclonal phenotypes. Expressed mutations seeding these subtrees can be evaluated by `Trisicell-PartF` to identify high confidence mutations strongly associated with each subclone. ^*a*^Alternative trees can also be obtained through other means of sequencing (e.g. DNA).

We assessed `Trisicell` on both simulated and real tumor sequencing data. In particular, we applied `Trisicell` to study melanoma, the malignant disease of melanocytes.

During embryonic development, neural crest (NC) differentiates to melanoblasts; they migrate and mature to melanocytes in skin. In normal skin, melanocytes accumulate mutations induced by ultraviolet radiation (UV) or other carcinogens over time. In rare instances, some melanocytes are transformed into melanoma. It was recently shown that human melanoma can be categorized into four distinct developmental subtypes [31] that are associated with expression of specific marker genes: “undifferentiated” subtype with the highest expression of *Axl*, “NC-like” subtype with high level of *Axl* and *Erbb3*, as well as “transitory” and “melanocytic” subtypes with lower level of *Erbb3* but high level of *Mitf*. In a recent study, we have also developed a “melanocytic plasticity signature” (MPS) score, which involves the expression levels of 45 genes: a higher MPS score correlates with higher plasticity and lower differentiation [32]. Importantly, therapeutic response to immune checkpoint blockade (ICB) is associated with the MPS score in both mouse and human melanomas.

For our recent study, we have generated multiple mouse melanoma models using distinct and relevant genetic alleles and carcinogenic procedures. Among them, the “M4” mouse melanoma model represents UV-induced RAS-mutated, differentiation-transitory subtype of human melanoma. This model responds to anti-CTLA-4 in a diverse manner, replicating the clinical treatment of melanoma with immune checkpoint inhibitors. Histological analysis of the tumors from this model also demonstrated a heterogeneous distribution of melanocytic makers’ expression values (*TRP2*, *TYRP1*, and *SOX10*), as well as pigmentation [32]. Therefore, it is a suitable model for studying intratumor heterogeneity.

In this study, we have applied `Trisicell` to sequencing data derived from the melanoma line B2905 derived from the M4 melanoma model. Specifically for this paper, we isolated and expanded 24 single cells from B2905 cell line to distinct “clonal sublines” that possess genomically uniform populations. Three tasks were performed: (i) On each subline, we performed high coverage bulk whole exome sequencing (bWES). (ii) We selected ~ 8 single cells from each subline and applied full-length transcriptome sequencing via the Smart-seq2 protocol [33] (scRNAseq). (iii) We finally selected 11 of these 24 sublines and implanted them in syngeneic C57BL/6 mice, each with 3 replicates, and performed bulk whole transcriptome sequencing (bWTS) on the resulting tumors (see Section 2.3 for the selection criteria).

The high coverage bWES data forms a “gold standard” for mutation calling in each of the 24 sublines. For each subline, we compared the mutations identified in bWES data to those derived from bWTS and scRNAseq data to identify those mutations that are expressed. Additionally, we evaluated “mutation calls” on both bWTS and scRNAseq data that are not present in bWES data to assess the scale of sequencing artifacts and RNA editing events [34] that lead to false positive mutation calls in transcriptome sequencing data. Since for some sublines we obtained bWTS data from three distinct replicates and scRNAseq data from (up to) eight single cells for each subline, these mutation calls are very robust. As a result, this tri-modal sequencing dataset offers a powerful way to benchmark computational methods for reconstruction and/or evaluation of mutation/clonal trees, especially through whole single-cell transcriptome sequencing.

We first employed simulated data to benchmark `Trisicell-Boost` using two existing methods for inferring mutation trees, namely SCITE and PhISCS, which we ran both as stand-alone programs as well as after boosting them with `Trisicell-Boost`. We observed that the run time performance of PhISCS as well as both the running time and accuracy of SCITE were substantially improved through the use of `Trisicell-Boost`, particularly when solving large problem instances.

Next, we assessed `Trisicell` on the tri-modal sequencing dataset generated from 24 clonal sublines of parental (B2905) cell line derived from the “M4” mouse melanoma model [32]. Having assessed that `Trisicell-Boost` (boosting, e.g. SCITE) achieves high levels of accuracy on simulated genomic data, we first applied it to the bWES data. The resulting tree clusters the most rapidly growing sublines in vivo in a distinct clade. The probabilities of specific mutations seeding this clade, as computed by `Trisicell-PartF`, is also very high, offering further evidence for the correctness of not only the topology of the inferred tree but also its mutational placements.

We then built trees for both bWTS data from 11 sublines (selected out of 24, as described in Section 2.3) and the scRNAseq data from all 24 sublines. `Trisicell-Cons` demonstrated that these trees were in strong agreement with what we obtained for bWES data. Importantly, these trees exhibited (almost) perfect clustering across cells (up to ~ 8 cells per subline, 175 cells in total) as well as replicates (3 replicates per subline) originating from the same subline, further validating the implied ancestor/descendant relationships between cells and sublines inferred through scRNAseq and bWTS data. Finally we compared the mutated genes specific to each of the four major clades we observed in the bWTS tree to those in human melanomas with distinct responses to ipilimumab (anti-CTLA4) therapy [35] and observed a striking similarity between the most responsive clades of the bWTS tree and the responder type human melanomas, suggesting convergent subclonal expansion across mouse and human tumors.

After extensively evaluating `Trisicell` on our tri-modal sequencing dataset, we applied it to a larger, preclinical, dataset involving B2905 melanoma cell line implanted in syngeneic (C57BL/6) mice, this time treated with anti-CTLA-4 immune checkpoint inhibitor. We sequenced a total of 764 cells from four anti-CTLA-4-treated tumors as well as two control tumors, via Smart-seq2 protocol, which, again, allowed the identification of mutations in full-length transcripts. The tree that we built for this large scale scRNAseq dataset features distinct clades strongly associated with differentiation status and therapeutic response, leading to the conclusion that a subset of seed mutations drove reshaping of the developmental landscape of the tumor, resulting in substantial phenotypic heterogeneity. Interestingly, the mutation profile of one of these clades overlapped with a major clade observed in the mutation tree of a patient-derived xenograft (PDX), further supporting convergent subclonal expansion between mouse and human melanomas.

## 2 Results

### 2.1 Development of Trisicell

`Trisicell` is comprised of three computational methods of independent use and interest: respectively for *assessing, integrating*, and *boosting* methods for reconstructing mutation trees so as to obtain reliable histories of tumor progression in a scalable manner. Importantly, we also present matching whole transcriptome and exome sequencing data from clonal sublines, each derived from a single cell of a melanoma mouse model, as well as single-cell transcriptome sequencing data from these sublines. As will be detailed below, by jointly evaluating `Trisicell` inferred and validated trees on these clonal sublines with matching gene expression data, we demonstrate that certain subpopulations of the melanoma mouse model seeded by specific mutations share developmental status, implying a connection to tumor progression. We expect that this dataset will help future method development efforts by offering a high-quality platform for benchmarking, parameter tuning, and integration with other methods.

`Trisicell` is comprised of the following three tools:

1. Available methods for inferring mutation trees of tumor progression from single-cell data aim to reconstruct the “most likely” tree using a noisy genotype matrix as their primary input. Even if all of the mutation calls in the genotype matrix are correct and the method used can guarantee optimality, the output tree is just one of the many possible interpretations of the input data as it only represents the “global” optimum. However, in many cases, we are interested in the likelihood (and the reliability) of “local structures”. This is commonly encountered in structural biology, e.g. in RNA-structure prediction where the goal is typically computing the most likely/thermodynamically stable secondary structure [36]. The secondary structure of an RNA strand can be represented as a tree and the likelihood of distinct subtrees of the secondary structure can be assessed through a “partition function algorithm” [37, 38].^3^ `Trisicell-PartF` features the first partition function formulation for a mutation tree of tumor progression and employs a sampling strategy to compute it for any user-defined subtree/subclone seeded by one or more mutations. Importantly, we prove that, as the sample size increases, the output of `Trisicell-PartF` converges to the correct value of the partition function.
2. In order to “solve” large instances of tumor progression reconstruction problem, available approaches make implicit and explicit assumptions on data behavior and, as a result, may output different trees for a given input genotype matrix. In fact, even the set of mutations themselves, and thus the genotype matrix derived from a read collection, may vary significantly as a consequence of the employed mutation calling pipelines. Furthermore, the type of sequencing data used for obtaining mutation calls (e.g., targeted or whole, genome or transcriptome sequencing) could result in distinct mutation calls for a given set of cells. As a result, it is many times not feasible to compare trees describing the lineages of a given set of cells from a tumor sample through their “mutation tree” representation, where each node is labeled with one or more distinct mutations. Unfortunately, since available measures for comparing mutation trees were typically not developed for single-cell data, they focus on how mutations (and not cells) are placed on distinct lineages.^4^ As will be described below, our `Trisicell-Cons` features a novel measure to compare two or more *cell lineage trees*. Here we use the broad definition of a cell lineage tree as a tree where leaves and some of the internal nodes are each labeled with one or more cells (in contrast to a narrower definition where only leaves are labeled with a unique identifier [23]). Our measure is based on a specific notion of a “consensus” between two trees, accompanied with an algorithm named `Trisicell-Cons` and its implementation to build this consensus. On a given set of cell lineage trees, `Trisicell-Cons` minimizes the number of edges to be “collapsed” so that the resulting trees are isomorphic. As such, `Trisicell-Cons` generalizes the consensus taxonomic tree construction algorithm of Day [43] for dendrograms, i.e. trees where only leaves are labeled with a single unique label. In order to modify Day’s algorithm for trees where any internal or leaf node may have an arbitrary number of labels^5^, we developed a new combinatorial approach and implemented it in `Trisicell-Cons`. We note that `Trisicell-Cons` provably obtains the largest possible consensus tree between the two input cell lineage trees in time quadratic with the input size. As we will demonstrate, it can be employed to integrate trees obtained by various methods and data types to help infer a more reliable history of a tumor progression.
3. `Trisicell-Boost` is a “booster” strategy for scaling up available methods for inferring mutation trees of tumor progression, especially those based on principled but relatively slow combinatorial approaches such as SCITE and PhISCS. It relies on the observation that the “correct” tree involving any subset of mutations in a tumor preserves the relative order of mutations with respect to their ancestor-descendant and different-lineage dependencies compared to the “correct” tree on the entire set of mutations. In its implementation, `Trisicell-Boost` first performs a sampling and tree inference step in which a specified number of mutations and cells from the input genotype matrix are sampled and, using the resulting submatrix as input, a mutation tree is built through any user-selected tool that works well on small-scale data. This atomic step is repeated for a user-defined number of times resulting in a collection of smaller trees on distinct subsets of mutations. Since these trees are obtained independently from each other, their inference tasks could be executed in parallel. Once this step is completed `Trisicell-Boost` reconciles them into a single mutation tree featuring the complete set of mutations (and cells).

An overview of `Trisicell` methods is depicted in Figure 1.

### 2.2 Benchmarking of methods for reconstructing mutation trees of tumor progression on simulated data

Using the procedure described in Supplementary Section I, which has been employed to assess many of the available methods for inferring mutation trees (including SCITE, PhISCS, OncoNEM, and B-SCITE), we simulated 80 distinct genotype matrices under infinite sites model, with a varying number of mutations (300 and 1000), cells (300 and 1000) and false negative mutation calling rates (0.05 and 0.20). The false positive mutation calling rate was set to 0.001 and the size of the simulated tree was set to 100 in all simulations (see Supplementary Section I for a more detailed discussion on the tree size and other parameter settings). ^6^

On each simulated dataset, we compared `Trisicell-Boost` against both SCITE and PhISCS as stand-alone programs.^7^ The accuracy of each tool for each parameter setting is provided in Figure 2. As can be seen, the results obtained by all three methods are comparable when n = m = 300. This is also the case when n = 1000 and m = 300 when false negative rate is 0.05, however PhISCS fails to report solutions for some instances when false negative rate is 0.2. When m = 1000 the performance of SCITE (with a 24-hour time limit) under both measures falls substantially below that of `Trisicell-Boost`. PhISCS also has difficulty handling certain problem instances in 24 hours when m = 1000, with the exception for n = 300 and the false negative rate of 0.05; for this setting it performs similar to `Trisicell-Boost`. It is worth noting that, with respect to different-lineage accuracy measure, `Trisicell-Boost` is near perfect across all simulations. With respect to ancestor-descendant accuracy measure, Trisicell-Boost also achieves very high accuracy, which increases as the number of simulated single cells is increased. An evaluation of the robustness of Trisicell-Boost to the presence of deletion events, which are a major cause of violations of infinite sites assumption, and additional information of running simulations can be found in Supplementary Section L.

**Figure 2.**
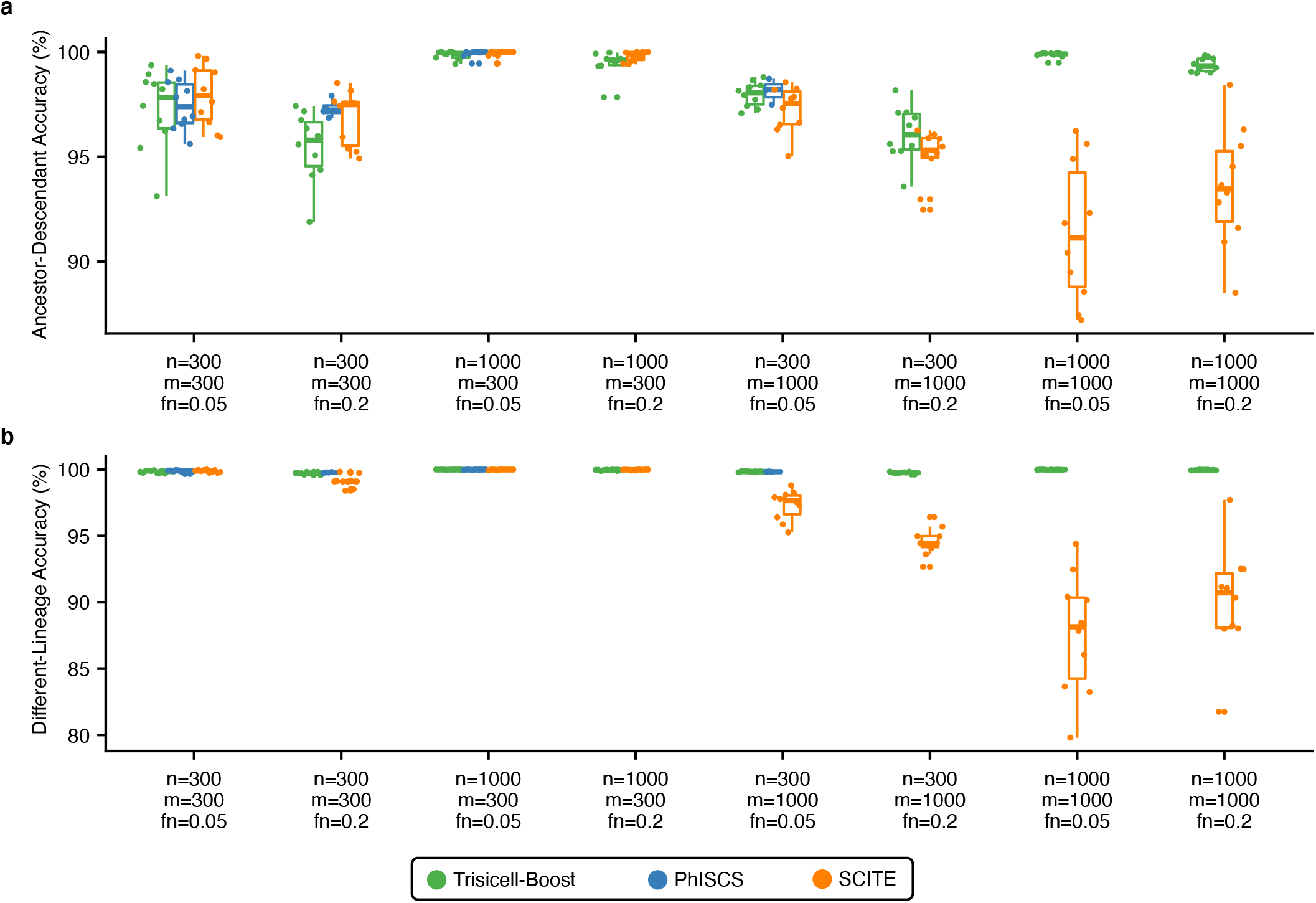
Benchmarking Trisicell-Boost on simulated data. A Comparison of SCITE and PhISCS with their `Trisicell-Boost` boosted version^*a*^ on simulated data, with respect to two commonly used measures of accuracy (see Supplementary Section J for definitions). Boosting SCITE and PhISCS does not result in loss of accuracy on smaller datasets that they can handle well as stand-alone tools. However, their performance is substantially improved by the boost they received from `Trisicell-Boost` on larger datasets. Details on the simulations and accuracy measures are provided in Supplementary Sections I and J. The parameters *n*, *m* and *fn* respectively denote the number of simulated single cells, mutations and the false negative rate in single-cell sequencing data. For each value of *m* (300 or 1000), 10 distinct trees of tumor progression were simulated and then for each value of *n* (300 or 1000) and *fn* (0.05 or 0.20) genotype matrices were derived from these trees. In all experiments, the false positive rate was set to 0.001 (i.e., 0.1%) and the missing entries rate was set to 0.05 (i.e., 5%). Note that PhISCS could not terminate within the 24-hour time limit on some instances of the problem - these cases are excluded from the results. The exact number of these instances can be found in Figure S17a. ^*a*^Even though SCITE and PhISCS use different combinatorial approaches, they have the same objective and constraints and thus their output on smaller trees (i.e. those which are employed by `Trisicell-Boost`) are isomorphic. As a result, we do not report on boosted versions of SCITE and PhISCS separately.

### 2.3 Benchmarking of methods for reconstructing mutation trees of tumor progression on clonal sublines from the M4 melanoma model

Melanocytes are derived from the neural crest during embryonic development. In normal skin, melanocytes accumulate mutations induced by ultraviolet radiation (UV) or other carcinogens over time. In rare instances, some of them are transformed into melanoma. It was recently shown that human melanoma can be categorized into four distinct developmental subtypes. Ranked with respect to differentiation levels, they are “undifferentiated”, “neural crest-like”, “transitory”, and “melanocytic” [31]. In a previous study [32], we have generated multiple mouse melanoma models using distinct genetic alleles and carcinogenic procedures with the aim of better modeling human melanoma. Previously, through comparative analysis, we have also identified a “melanocytic plasticity signature” (MPS) composed of 45 genes, which allows quantification of the developmental status of melanoma: a higher MPS score correlates with higher plasticity and lower differentiation [32]. We have found that therapeutic responses to immune checkpoint inhibitors are associated with MPS scores in both mouse and human melanomas. In this study, we used the B2905 cell line derived from “M4” mouse melanoma model that represents UV-induced RAS-mutated, differentiation-transitory subtype of human melanoma. This model responds to anti-CTLA-4 in a diverse manner [32], replicating the clinical treatment of melanoma with immune checkpoint inhibitors. Therefore, it is a suitable model for studying tumor heterogeneity.

In order to investigate `Trisicell-Boost`’s applicability to experimental datasets, we generated three types of sequencing data from tumors derived from the parental M4 (B2905) mouse melanoma model. First, 24 single cells from M4 were expanded into clonal sublines, and then subjected to bulk whole exome sequencing (bWES). We then implanted each subline into five syngeneic mice to monitor their growth kinetics. Out of the 24 sublines, 11 reached 100% tumor-take rate. Each of these tumors was harvested at an endpoint, and 3 tumors from each subline were subjected to bulk whole transcriptome sequencing (bWTS) as triplicates (Figure 3a). Additionally, we extracted 8 single cells from each subline and subjected them to Smart-seq2 protocol to obtain scRNAseq data (Figure 4a).

**Figure 3.**
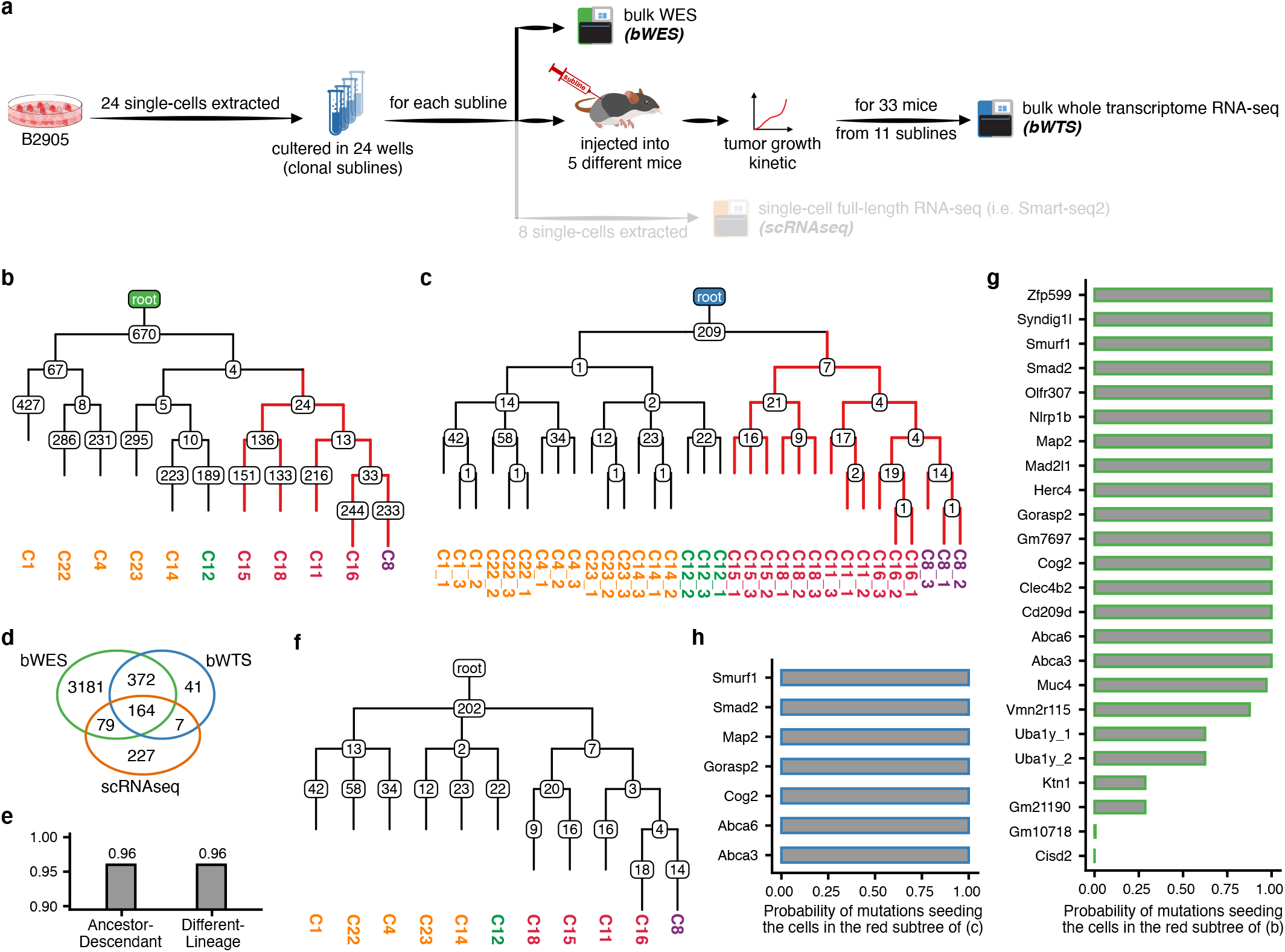
Comparison of trees of tumor progression inferred by Trisicell on bulk whole transcriptome sequencing (bWTS) vs bulk whole exome sequencing (bWES) data from 11 clonal sublines.*^a^*, (a) Overview of the experimental protocol specific to bWES and bWTS data generation. (b) bWES-based mutation tree obtained by `Trisicell-Boost`. The number of somatic de novo mutations is shown for each node. The color used for the label of each subline indicates its “aggressiveness”. In decreasing order of aggressiveness they are: red, orange, purple, and green.*^b^* (c) bWTS-based mutation tree obtained by Trisicell-Boost; each subline had five independently grown replicates; the fastest growing three (see Figure S4) were sequenced. (d) The number of shared and private “mutation” calls across bWES and bWTS, as well as scRNAseq data (see Figure 4 for results on scRNAseq data). (e) Commonly used measures of accuracy for the bWTS tree with respect to the bWES tree (see Supplementary Section J for definitions). (f) `Trisicell-Cons` inferred consensus tree for bWES and bWTS trees. The number of somatic de novo mutations shown in the nodes of the consensus tree are for those shared between the bWES and bWTS trees. (g) `Trisicell-PartF` calculated the probability of each mutation seeding the aggressive subtree of the bWES tree, comprised of sublines C15, C18, C11, C16, and C8.*^c^* (h) `Trisicell-PartF` calculated the probability of each expressed mutation seeding the subtree comprised of the most rapidly growing tumors in the bWTS tree.*^d^* *^a^*See Figure S4 for the criteria used to select these 11 sublines out of 24. ^*b*^Aggressiveness of a subline is measured with respect to the median survival time (in days) and grouped by quartiles (red: <Q25, orange: Q25-median, purple: median-Q75, green: >Q75). ^*c*^A mutation with a high probability is likely to have occurred at the root of a subtree that is comprised of the sublines C15, C18, C11, C16, and C8. ^*d*^All expressed mutations have probability 1 but some that are not expressed have lower probabilities.

**Figure 4.**
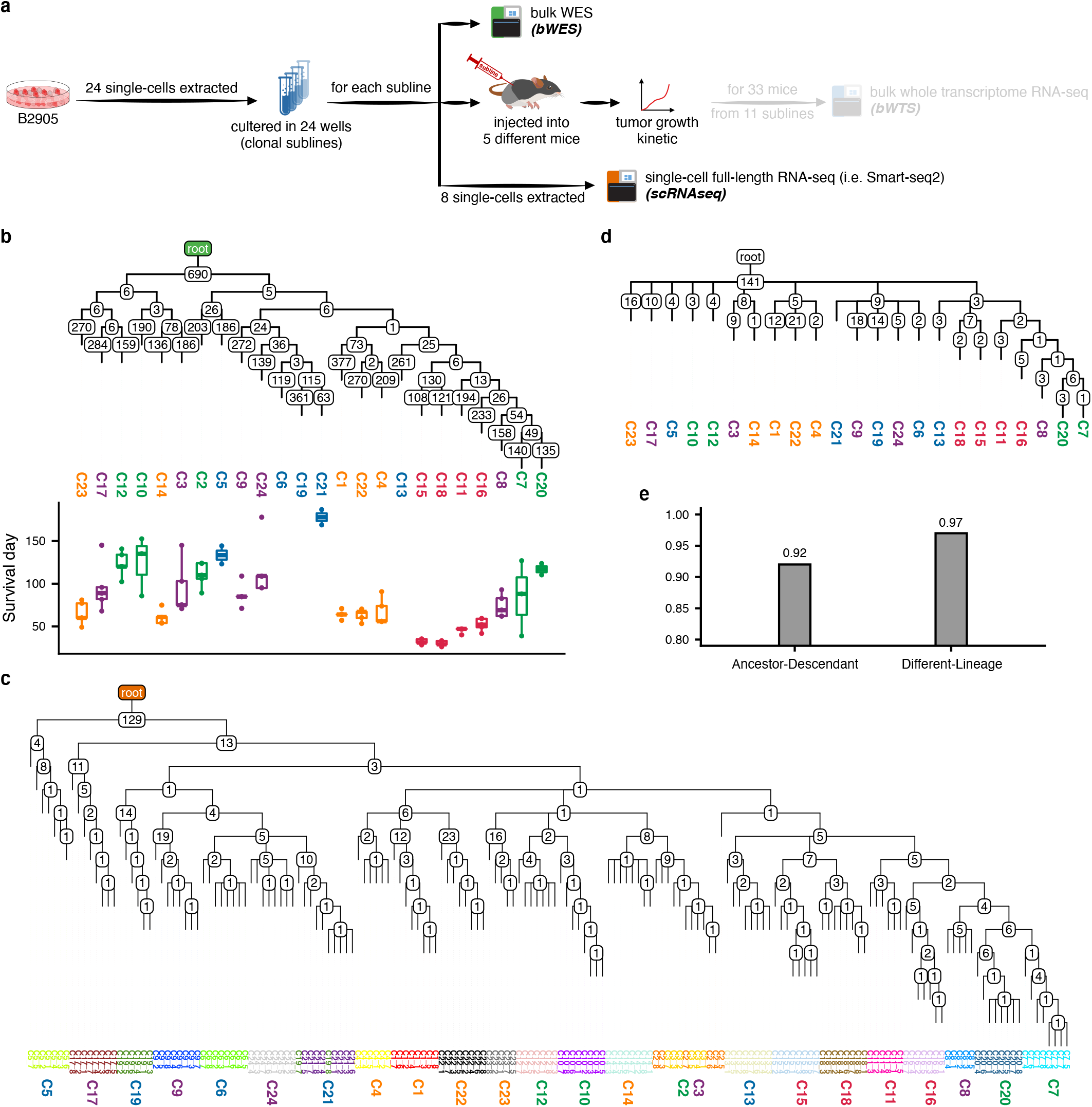
Comparison of trees of tumor progression inferred by Trisicell on single-cell RNA-seq (scRNAseq) vs bulk whole exome sequencing (bWES) data from 24 clonal sublines. (a) Overview of the experimental protocol specific to bWES and scRNAseq data generation. (b) bWES-based mutation tree inferred by `Trisicell-Boost`. The number of somatic de novo mutations is shown for each node. The color used for the label of each subline indicates its aggressiveness. In decreasing order of aggressiveness they are: red, orange, purple, green, and blue.*^a^* (c) scRNAseq-based mutation tree inferred by `Trisicell-Boost`; from each subline ~ 8 cells were sampled and sequenced. Cell colors indicate the subline they were sampled from. The cells for each subline are well clustered, with the exception of C2 and C3, which are mixed.*^b^* (d) `Trisicell-Cons` inferred consensus tree for bWES and scRNAseq trees. The number of somatic de novo mutations shown in the nodes of the consensus tree are for those shared between the bWES and scRNAseq trees. (e) Commonly used measures of accuracy for the scRNAseq tree with respect to the bWES tree (see Supplementary Section J for definitions). ^*a*^Aggressiveness of a subline is measured with respect to the median survival time (in days) and grouped by quartiles (red: <Q25, orange: Q25-median, purple: median-Q75, green: >Q75, blue: <3 tumors formed). ^*b*^In the consensus tree of panel (d), only C3 is represented. Double missense mutations + splice mutations

For assessing `Trisicell` on this tri-modal sequencing dataset, we first built the tree of the 24 clonal sublines, through mutation calls that we made on their bWES data using `Trisicell-Boost` (Figure 4b). Next, we built the tree of the 11 sublines (each with 3 replicates) through mutation calls that we made on their bWTS data (Figure 3c). Then, by the use of `Trisicell-Cons`, we constructed the consensus of the two trees that we had built through the use of bWES data and bWTS data, which exhibited only minor differences (in the location of the subtree with sublines C1, C4, and C22; see Figure 3f). Furthermore, the tree we built using bWTS data was very similar to that using bWES data with respect to commonly used measures of accuracy (Figure 3e). Note that, even though the exact set of mutation calls made on any given subline from bWTS data differed from that of bWES data, the mutation calls made on the bWTS data differed minimally across the three replicates of the same subline (see Figure 3c for those mutations also observed in bWES data and Figure S19 for the entire set of mutations). As a result, the `Trisicell-Boost` built mutation tree on bWTS data was able to cluster all replicates from each subline correctly, demonstrating its potential use on mutations derived from RNAseq data for reconstructing history of tumor progression.^8^ In contrast, DENDRO and Cardelino, the only other tools for tumor progression history inference through RNAseq reads, both failed to cluster replicates of certain sublines correctly, as discussed in more detail below.

The mutation tree built on bWES data features a subtree comprised of the most rapidly growing sublines (measured by the survival time^9^) in vivo (Figure 4b). These sublines, i.e., C11, C15, C16, and C18, are seeded by a distinct set of mutations, several of which are also observed in bWTS data.^10^ An analysis of this subtree in both bWES and bWTS trees by Trisicell-PartF (see Figure 3g,h) suggests that some of these seeding mutations (especially the seven observed in both bWTS and bWES data, all with high probability) may have contributed to the emergence and proliferation of the associated subclone. In particular, *Map2* is a known melanoma cell proliferation inhibitor [44]. A non-synonymous SNV in exon9:c.G739C:p.G247R of this gene is likely one of the contributors to the rapid tumor growth in these sublines.

As mentioned earlier, we have also used our bWTS dataset to evaluate DENDRO and Cardelino, the only other tools for inferring the history of tumor progression via RNAseq reads, in addition to SCITE and PhISCS, this time as stand-alone tools. DENDRO produced a tree which is substantially different from the bWES tree (e.g. it clustered the most aggressive sublines with some of the least aggressive); in fact, it even failed to cluster replicates of certain sublines correctly. Cardelino on the other hand is primarily developed for assigning mutations to the nodes of a preset topology and could not infer trees of this scale - see Supplementary Sections Q and R). SCITE and PhISCS on the other hand produce trees (almost) isomorphic to their boosted versions with `Trisicell-Boost`, providing further evidence for the robustness of `Trisicell-Boost` in scaling up these methods; see Supplementary Section P. (Note that for brevity we only provide the results for SCITE as PhISCS results were nearly identical.)

We finally built the scRNAseq based mutation tree of ~ 8 single cells derived from each of the 24 sublines. A total of 175 (out of 24 × 8 = 192) cells provided sufficient read coverage to perform mutation calling. In order to reduce transcriptome sequencing artifacts and RNA editing, we only consider those mutations shared by the corresponding bWES data. Finally, to account for deletion events (leading to loss of heterozygosity LOH), if a mutation in a gene with a copy number of 1 is not observed in a cell, we set its status to “NA” (i.e. unknown) (see Supplementary Section S for details of copy number profiling). The resulting tree obtained by `Trisicell-Boost` (again boosting SCITE) is provided in Figure 4c. Notice that the cells originating from the same subline are well clustered, akin to the clustering of replicates from each subline in the bWTS tree. Furthermore, the major subtrees of the scRNAseq tree are identical to that of the bWES tree as demonstrated by their `Trisicell-Cons` consensus shown in Figure 4d. For example, the subtree featuring the most rapidly growing sublines in vivo (i.e. C11, C15, C16, C18) in bWES tree is preserved in the scRNAseq tree. Additionally, ancestor-descendant and different-lineage mutation pair relationships in the scRNAseq tree with respect to the bWES tree are highly preserved (Figure 4e). Overall, the scRNAseq tree demonstrates that `Trisicell-Boost` can be used to infer detailed and scalable trees of tumor progression through the use of scRNAseq data, especially when combined with bulk whole genome or exome sequencing data for mutation calling. (The corresponding mutation trees that we built for all “mutations” observed in bWTS and scRNAseq data can be found in Supplementary Section O.)

We also evaluated DENDRO and Cardelino, as well as SCITE and PhISCS as stand-alone tools, on the scRNAseq dataset. As in the case of some larger simulations, PhISCS could not complete the task in 24 hours; this is because it necessarily searches for the minimum possible noise reduction for obtaining a mutation tree. SCITE on the other hand produced a result since it is designed to search for a solution in a user-specified time, but its output could be sub-optimal. The tree built by SCITE is given in Figure S22: even though SCITE clusters cells from each subline well, the relative placement of the sublines does not agree with that in the bWES tree as demonstrated by their `Trisicell-Cons` built consensus. DENDRO, in contrast, failed to cluster cells from the same subline Figure S24. Similarly, Cardelino, even when used for cell clustering only, fails to perform well on this dataset (see Supplementary Section R).

### 2.4 Overlap between the mutated gene profiles of sublines and human melanoma patients treated with anti-CTLA-4 immune checkpoint inhibitor

To explore the clinical significance of the trees of tumor progression established by Trisicell, we investigated the 11 clonal B2905 sublines whose in vivo tumors were subjected to bWTS, and compared them to bWTS data from human melanomas. Van Allen et al have analyzed pre-treatment melanomas of patients receiving ipilimumab (anti-CTLA4 immune checkpoint blockade) treatment [35] and categorized the patients into three response groups: no/minimal clinical benefit (non-responders, NR), long-term survival with no clinical benefit (long-term survivors, LS), and clinical benefit (responders, R). The analysis of the bWTS data showed that, although majority of the samples were from NR patients (56%), and only 32% belonged to R patients (Figure 5a, upper panel), R melanomas contributed far more mutated genes (47% vs. 32%, p-value = 0.0419) and distinct mutations (50% vs. 30%, p-value = 0.0410) than the NR melanomas in all samples (Figure 5a, middle and lower panels). Taken together, these indicate that higher number of expressed mutations are associated with more favorable response to ipilimumab anti-CTLA4 treatment.

**Figure 5.**
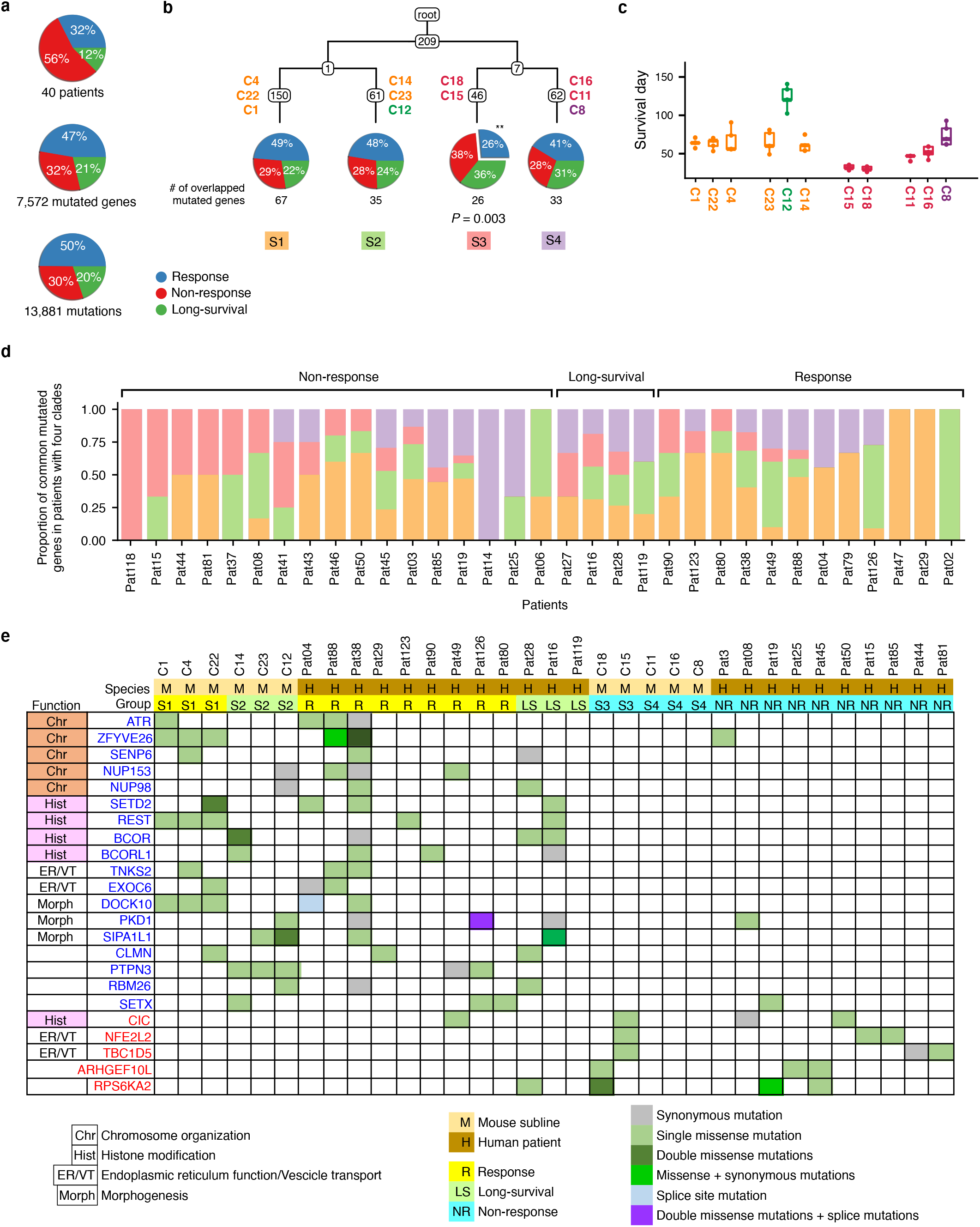
Overlap between the mutated gene profiles of 11 sublines Figure 3c and (human) melanoma patients treated with anti-CTLA-4 immune checkpoint inhibitor (ipilimumab) [35]. A total of 40 metastatic human melanoma patients were stratified into three response groups based on RECIST criteria [59], duration of overall survival and duration of progression-free survival: response (benefited therapy) - depicted in blue, non-response (no benefit from therapy) - depicted in red, and long(er) survival (early tumor progression, thus limited benefit from therapy but longer survival) - depicted in green. (a) Top pie chart: distribution of the 40 patients across three response groups. Middle pie chart: The distribution of mutated genes across 40 patients in the three response groups (The number of genes mutated per patient are summed up for each patient group). Bottom pie chart: The distribution of mutations across the 40 patients in the three response groups. (The number of mutations across all genes per patient are summed up for each patient group). As can be seen, even though the cohort is comprised of mostly non-responders, half of the mutations are observed in responders, suggesting that several are likely passenger mutations. (b) Four major clades of the tumor lineage tree from Figure 3c: S1 (comprised of mouse melanoma sublines C4, C22, C1, all with longer survival), S2 (comprised of C14, C23, C12, all with longer survival), S3 (comprised of C18 and C15 with the shortest survival), and C4 (comprised of C16, C11 and C8, all with mid-range survival). The number of (expressed) mutated genes in each clade which are also mutated in at least one patient in each response group is depicted in the pie charts. Even though many of the mutations observed in human melanomas are likely passengers, the mutated genes specific to the most aggressive clade S3 are less commonly mutated in the responder group (only 26% of mutations in S3 are shared with responders, whereas 47% of the mutated genes are in the responder group, p-value = 0.003). A similar (but less strong) association can be observed for the (less aggressive) clade S4. In contrast, the distribution of (expressed) mutated genes in the least aggressive clades S1 and S2 (which, again, overlap with the human melanomas) closely match their distribution across the three response groups in panel (a). (c) Survival time distribution of the sublines (each obtained from 5 mice); the sublines are sorted with respect to their placement in the tumor lineage tree. (d) The proportion of mutated genes among each patient in the cohort shared with the four clades S1 (orange), S2 (green), S3 (red) and S4 (purple) of the cell lineage tree from (b). Out of 40 patients in the cohort, 8 had no common mutated genes with these four clades and thus are not depicted here. Non-responder patients have a higher proportion of mutated genes that are specific to the more aggressive clades of the tumor lineage tree. In fact each responder’s mutated gene profile overlaps more (≥ 50%) with the less aggressive clades (S1 and S2) while that of 10 out of the 17 non-responders and 2 out of the 4 long-survivors overlap more (≥ 50%) with more aggressive clades (p-value = 0.0017). (e) Genes recurrently mutated in human melanomas as well as in at least one tumor of the four clades, along with gene function and mutation type information. Those (typically singly missense) mutations observed in genes involved in chromosome organization and morphogenesis are specific the less agressive S1 and S2 clades and the R type human melanomas.

The 11 clonal sublines of mouse melanoma were clustered by the `Trisicell-Cons` consensus tree to four major clades: S1 (formed by C1, C22, C4), S2 (formed by C23, C12, C14), S3 (formed by C15, C18), and S4 (formed by C11, C16, C8) (Figure 3). Interestingly we have observed significant differences in the growth kinetics of the in vivo tumors among these four clades. In general, tumors in S1 and S2 consistently reached the endpoint later than S3, whose tumors had the most rapid growth (see Figure 5c), and tumors of S4 grew at rates in-between these two extremes. To model human melanoma treated with ipilimumab, we selected one subline from each clade, specifically C1, C14, C15, and C11, to test their responses to anti-CTLA4. These four sublines exhibited diverse responses. The Mantel-Haenszel hazard ratio of anti-CTLA4 treatment vs. control in mice carrying C1, C14, C15, and C11 tumors was 0.096, 0.334, 0.169, and 0.405, respectively (see Supplementary Section G). However, the medium survival time of anti-CTLA4-treated mice carrying C1, C14, C15, and C11 tumors are “undefined” (i.e. 20% complete response), 65 days, 32 days, and 46 days, respectively. Therefore, C1 tumors can be defined as R, C11 and C15 as NR, and C14 as LS melanoma, following the classification by Van Allen et al. [35].

Intriguingly, the three R tumors in clade S1 also exhibited higher number of mutated genes than all tumors in the other three clades, similar to the association between R tumors and higher number of mutated genes in human melanomas (Figure 5a, middle panel). However, the number of mutated genes or mutations is not sufficient to explain the LS phenotype in the human or the mouse melanomas. This motivated us to investigate how the “quality” of the mutated genes impact the therapeutic response by comparing the sets of mutated genes in distinct anti-CTLA4 response groups across human and mouse melanomas. For that we identified the genes mutated in at least one mouse tumor as well as one human melanoma sample (“overlapped genes”), and then analyzed their distribution among the melanomas of both species. As shown in Figure 5b, the distribution of the overlapped genes in the least aggressive clades, S1 and S2 closely match the distribution of mutated genes across the three response groups in Figure 5a. In contrast, the mutated genes specific to the most rapidly growing clade S3 are significantly less commonly mutated in the human R melanomas (only 26% of mutated genes in S3 are shared with human R melanomas, which harbor 47% of the mutated genes across all patients; p-value = 0.003). A similar but weaker association can be observed for the more slowly growing tumors in clade S4.

We also examined the distribution of the overlapped mutated genes from clades S1 (orange), S2 (green), S3 (red) and S4 (purple) across human melanomas (Figure 5d). Seven of the human melanomas had a bWES sequencing depth of only ≤ 10% of the average sequence depth, and thus exhibited very few mutations; none of these mutations were observed in the overlapped genes (with these four clades) and thus these melanomas are not included in this figure. For the remaining 33 human melanomas, the number of overlapped genes are, on average, 2.8% of the total number of mutated genes (the proportion of overlapped genes among all mutated genes varied between 0.9% and 13.4% across the 33 melanomas and had no association with therapy response).

Interestingly, when examining the mouse clade composition of the overlapped genes, NR melanomas have a significantly higher proportion of mutated genes specific to S3 and S4, the more aggressive clades of the tumor lineage tree (p-value = 0.0068). In fact the mutated gene profile of each R (responder) melanoma overlaps more (≥ 50%) with the less aggressive clades (S1 and S2) while that of 10 out of the 17 NR (non-responder) and 2 out of the 4 LS (long-survivor) melanomas overlap more (≥ 50%) with the more aggressive clades (p-value = 0.0017). Taken together, these results suggest that the tumor progression tree identifies distinct subclones with specific mutation profiles, which have differential response to therapy, even though many of the mutations observed in human melanomas are likely passenger events. Importantly, the “convergence” of the mutated gene sets in mouse and human melanomas gives functional implication of these genes. In fact, overlapped genes that are recurrently mutated in melanomas of R patients are significantly enriched in chromosome organization (including ATR, ZFYVE26, SENP6, NUP153, NUP98) and DNA methylation functions (including SETD2, REST, BCOR, BCOR1). (see Figure 5e for a detailed presentation for commonly and recurrently mutated genes across melanomas, together with function and mutation type information) Recent studies have demonstrated that mutations in DNA repair genes [45] and alteration in histone modification [46] are associated with the efficacy of immune checkpoint blockade therapy.

### 2.5 Application of Trisicell to anti-CTLA-4-treated tumors derived from the M4 melanoma model

To investigate the origin of intratumoral heterogeneity and its impact on the resistance to anti-CTLA-4, we performed a preclinical study involving six mice implanted with M4 melanoma, four of which were treated with anti-CTLA-4 antibodies, and the remaining two with generic immunoglobulin IgGb as control (Figure 6a).^11^ A total of 764 cells from all six tumors that passed the quality control and had sufficient read depth were selected for downstream analysis.

**Figure 6.**
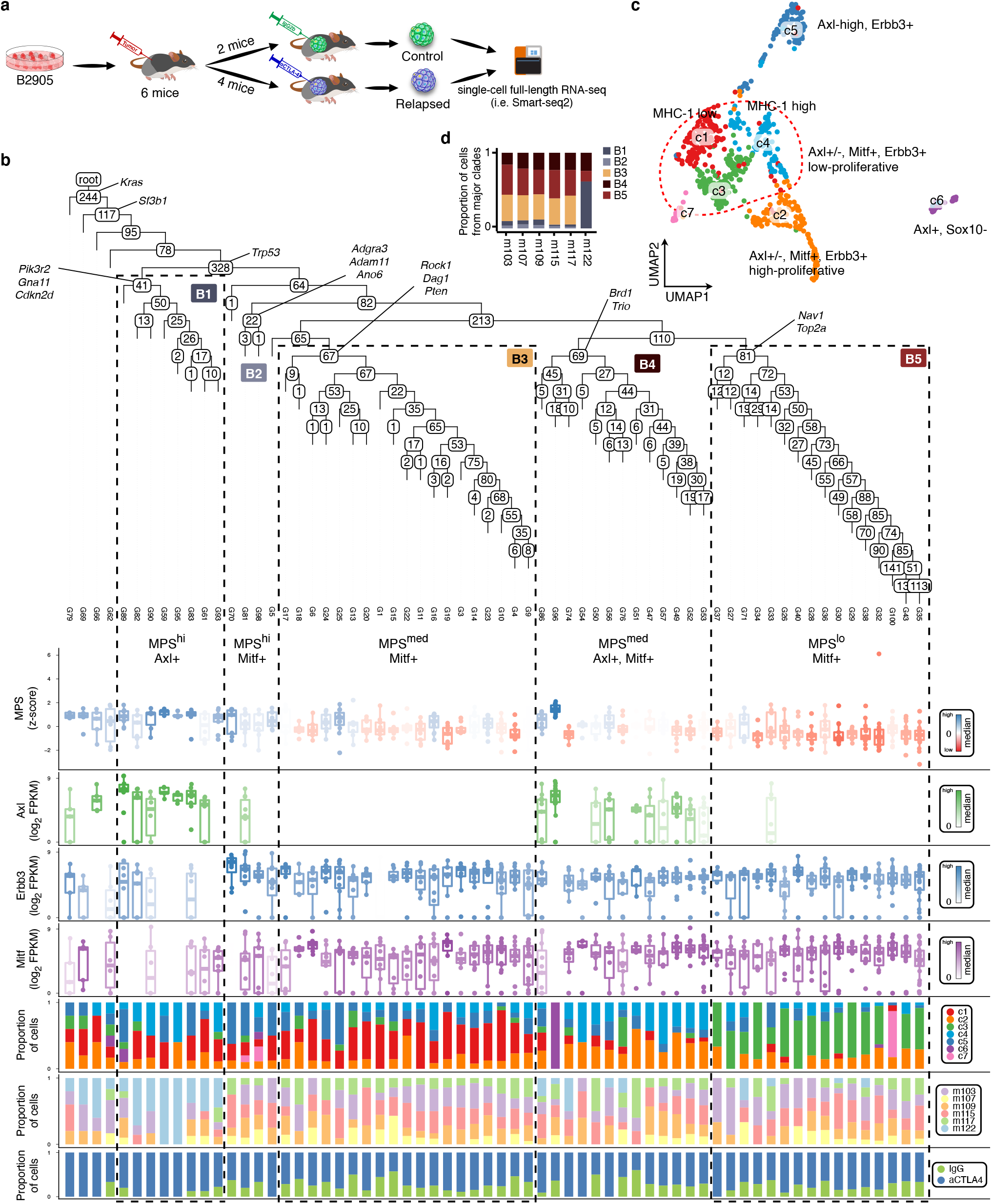
Mutation tree of tumor progression inferred by Trisicell-Boost on scRNAseq data from M4-derived tumors treated with anti-CTLA-4 immune checkpoint inhibitor. (a) Overview of the experimental protocol specific to this scRNAseq data. (b) `Trisicell-Boost` inferred mutation tree where each leaf represents a group of cells with similar mutational profiles. The number of somatic de novo mutations is shown for each node. For each leaf, (i) the corresponding melanocytic plasticity signature (MPS) score and the expression values of lineage markers, (ii) *Axl*, (iii) *Erbb3*, and (iv) *Mitf* are shown in the panels below the tree. Based on these, five major subtrees/clades, B1, B2, B3, B4, and B5 can be distinguished. Additional panels depict the proportion of cells in each leaf with respect to (v) expression profile based UMAP clustering, (vi) mouse id, and (vii) the type of treatment they received. (c) Clustering of cells with respect to overall gene expression profiles via UMAP (clusters are annotated by lineage markers). (d) The proportion of cells from each major subtree/clade of the inferred mutation tree in each mouse.

#### Trisicell analysis

To study the subclonal structure of the anti-CTLA-4 treated M4 melanoma, we applied `Trisicell` to reconstruct the mutation tree from all six tumors. The cells were first locally grouped into 64 mutationally homogeneous clusters and each cluster’s reads were re-evaluated to establish their mutation profiles (a similar approach for clustering cells prior to mutation calling have been used earlier, e.g. for CNV profiles [26, 47]). Trisicell-Boost inferred tree on this dataset is provided in Figure 6b.

As a next step, for each leaf node we calculated the median of (1) “melanocytic plasticity signature” (MPS) [32] and (2) expression of neural crest (NC) lineage developmental markers including *Axl* (undifferentiated/NC), *Erbb3* (NC/transitory), and *Mitf* (differentiated) [31]. To compare the phenotypes, we also analyzed the proportion of (1) cells from distinct UMAP clusters (see below for UMAP analysis); (2) cells from distinct tumors; and (3) cells from control vs anti-CTLA-4 treatment groups. When tracing specific mutations in each node, the “hotspot” mutations at *Kras*, *Sf3b1*, and *Trp53* (*p53*) were found in the five internal nodes closest to the root. This is expected since these are the well-known “driver” mutations and they frequently occur in human melanoma. Interestingly, the inferred mutation tree features five major subtrees (or clades), which are strongly associated with MPS values, as well as expression of *Axl* and *Mitf* : clade B1 (MPS^high^, Axl^+^), clade B2 (MPS^high^, Mitf^+^), clade B3 (MPS^medium^, Axl^−^, Mitf^+^), clade B4 (MPS^medium^, Axl^+^, Mitf^+^), and clade B5 (MPS^low^, Mitf^+^). Across these subtrees, the expression of *Erbb3* is similar to *Mitf*. The seeding mutations of these subtrees include those in *Pik3r2*, *Gna11*, *Adgr3*, *Pten*, *Brd1*, and *Top2a*, genes that are recurrently mutated in human melanoma.

The identification of distinct clades in the mutation tree allowed us to trace the subclonal tumor progression as a consequence of anti-CTLA-4 treatment. To examine the association between the tumors and clades, we calculate the proportion of cells from each tumor and treatment group for each leaf, as shown in the sixth and seventh panels under the tree in Figure 6b, respectively. While four clades (B2 to B5) do not enrich cells from specific tumors, B1 has significantly higher number of cells from tumor m122 of the anti-CTLA-4 treatment group. In correspondence, we investigated the subclonal response to the treatment by calculating the proportion of cells from each clade for each tumor; see Figure 6d. Observe that during the growth of tumor m122, clade B1 became predominant while B2 and B3 diminished, which suggests that these three clades were subjected to strong selection pressure. The fraction of clade B1 also slightly increased in tumor m103 and m109, but not in m115. Interestingly, our previous study had demonstrated that melanoma resistance is associated with high plasticity, consistent with the fact that B1 clade exhibits the highest MPS scores and expression of “undifferentiated” markers[32].

Our new results suggest that subclonal selection may not happen in every tumor responding to therapy. Since it has been reported that cancer cells can adapt to the therapeutic stress via a non-genetic mechanisms [48], we analyzed differentially expressed genes in each clade by comparing anti-CTLA-4 treated tumors against control tumors (see Supplementary Section E) and then subjected them to Gene Set Enrichment Analysis (GSEA) (see Supplementary Section F). Interestingly, differentially expressed genes of clade B1 are mostly enriched in inflammatory pathways, especially in m122, suggesting that cells of clade B1 were “intrinsically” resistant and thus selected by the immune response. This is consistent with our previous discovery that undifferentiated subtype of melanoma is resistant to immune checkpoint blockade [32], as clade B1 cells exhibit high MPS scores and Axl expression. Cells in other clades seem to activate pathways relevant to mesenchymal stem cells (myogenesis and cholesterol homeostasis in B2 and B3), mitotic arrest (mitotic spindle and G2M checkpoint in B4), and energy metabolism (oxidative phosphorylation in B5).

#### Expression profile based clustering of cells with UMAP

In order to compare tumor progression analysis with gene expression profile analysis we employed Seurat package [49]^12^ which resulted in seven clusters with distinct gene expression patterns (see Figure 6c), corresponding to differentially expressed gene sets (see Supplementary Section H). We examined the expression of lineage markers in these clusters and noticed that clusters c5 and c6 in Figure 6c have higher median MPS scores and more Axl^+^ cells than others, suggesting that they are less differentiated. Among these clusters c6 includes no Erbb3^+^ cells indicating that it is even less differentiated than c5. In contrast, the remaining clusters have lower MPS scores and more Mitf^+^ cells, indicating that they are more differentiated. Noticeably, the distribution of the expression patterns of these genes is heterogeneous, even among the cells from the same cluster. For example, Axl^+^ cells are also present in the five clusters with lower MPS scores. This implies that the UMAP clusters are only weakly associated with developmental status.

#### A comparison of Trisicell inferred mutation tree and UMAP clusters

We calculated the proportion of cells from each UMAP cluster (see Figure 6c) in each leaf node of the mutation tree. As can be seen in the fifth panel of Figure 6b, clade B5 is enriched with UMAP clusters c3 and c7, and clade B3 is enriched with UMAP cluster c1; all these clusters are Mitf^+^. In contrast, the two Axl^+^ clades, B1 and B4, are not enriched with cells from any particular UMAP cluster. These results indicate that the mutation-based clades in the tree are associated with distinct developmental status defined by MPS score and lineage markers, rather than the overall gene expression patterns as suggested by UMAP clusters.

In conclusion, our study demonstrates that Trisicell inferred mutation tree from transcriptomic mutational profiles of single cells can robustly trace distinct subclones, making it feasible to study their specific treatment responses. Our results are consistent with recent studies reporting on distinct subclonal populations that develop specific resistance or adaptation mechanisms to therapy [53, 54].

### 2.6 Application of Trisicell to single-cell sequencing data from human melanoma patient-derived xenograft

We applied Trisicell to analyze the single-cell RNA seq data of a melanoma patient-derived xenograft (PDX) in mice [55], generating the mutation tree shown in Figure 7, which includes mutations observed in the single cells obtained from tumors sampled at four time points following combined treatment of BRAF and MEK inhibitors. More than 70% of the mutations occurred in the trunk, and particularly in the first five nodes following root, indicating higher clonality. This may reflect the selection happening during in vitro culture and drug treatment of the PDX. The overall MITF and ERBB3 expression in the nodes of this tree is comparable to that of the B3 and B5 clades in the mouse M4 tree (Figure 6b) while AXL was expressed at a low level and inconsistently. Nevertheless, the two main subtrees in this PDX tree (namely “left” and “right” subtrees) are respectively associated with medium-to-high vs low MPS scores, indicating that they exhibited different developmental status, similar to the B3 and B5 clades in the mouse M4 tree (Figure 6).

**Figure 7.**
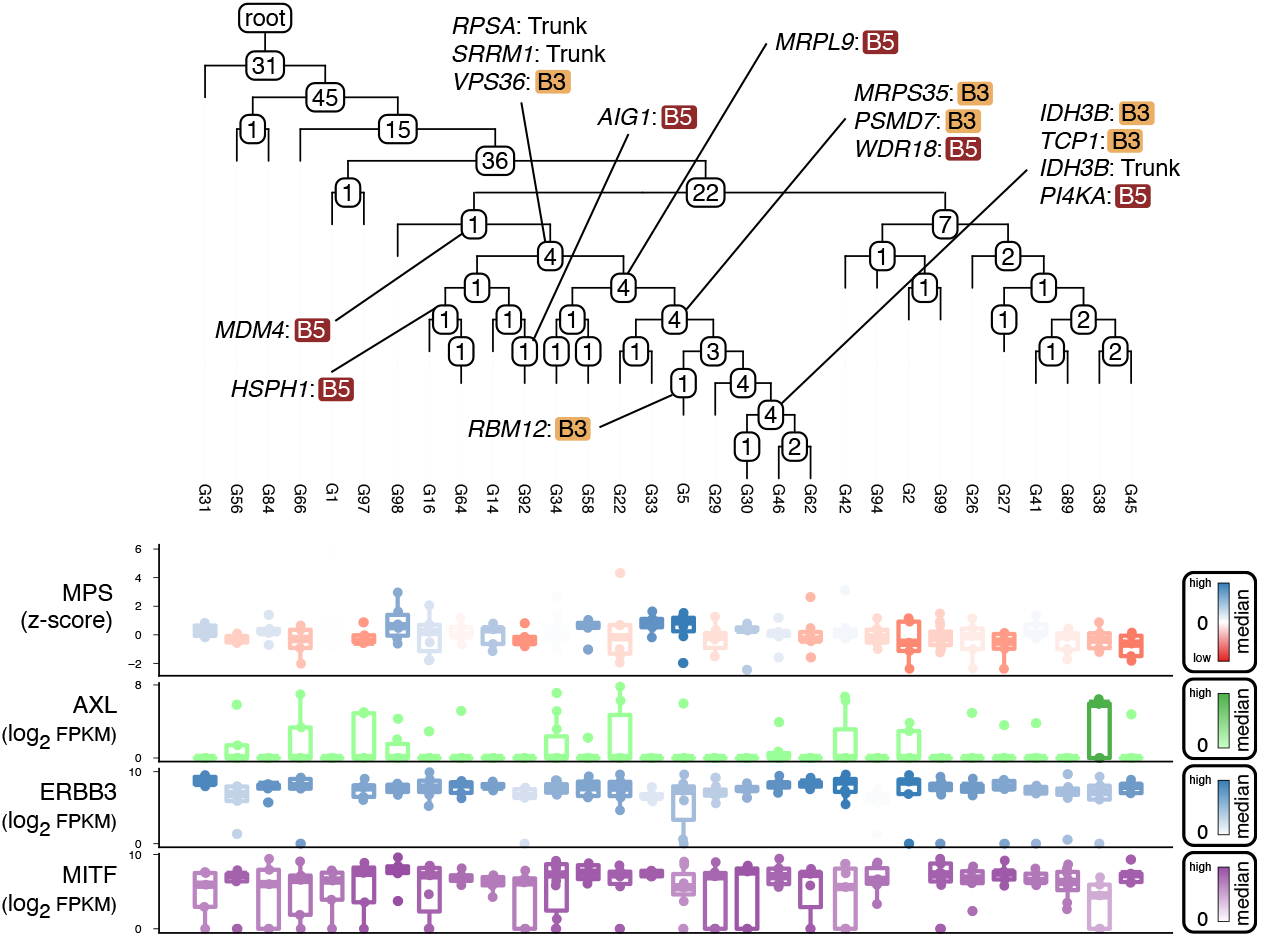
Mutation tree of tumor progression inferred by Trisicell-Boost on scRNAseq data from Minimal Residual Disease (MRD) sample. Human melanoma xenografts were treated with a BRAF and MEK inhibitors. Cancer cells were sampled at four distinct time points for scRNAseq [55] on which the tree was generated based on expressed mutations. Each leaf of the tree represents the pseudo-bulk mutational profile of a group of these cells with similar mutational composition (see [47]). The number of somatic de novo mutations occurring in each leaf or internal node is shown in the node. MPS score as well as the expression of three neural crest lineage developmental markers AXL, ERBB3, and MITF for each leaf are shown in the lower four panels. The mutation tree has two major subtrees. The mutated genes in the left-subtree which overlap with genes mutated in the mouse M4 tree (Figure 6b) are shown on the subtree, and are labeled with the distinct clade of the mouse M4 tree (B3 or B5) where they are mutated (the genes mutated in the first six nodes following the root of the mouse M4 tree are labeled as “Trunk”). While the earlier stages of the left-subtree overlap more with the Trunk or the B5 clade of the mouse M4 tree, the later stages show higher similarity to the B3 clade. Overall, the MPS (medium to high) as well as the expression of AXL (low expression), ERBB3 (high expression) and MITF (high expression) on the left-subtree is similar to the B3 clade of the mouse M4 tree.

Interestingly, out of the 197 genes with 206 distinct mutations in the PDX tree, 57 are also mutated in mouse M4 tree (28.9% with p-value < 0.005, see Figure 6). Among these 57 commonly mutated genes, 14 (IDH3B has two mutations) were mutated in the left subtree of the PDX tree; all but two of those (RPSA and SRRM1) were also mutated in clades B3 and B5 of mouse M4 tree (with p-value ≤ 0.002). Notably, similar to the PDX tree, B3 and B5 also have high MITF expression with negligible Axl expression. The association between mutated gene sets and developmental status across the subtrees of the two scRNAseq trees (of melanoma, derived from human PDX vs mouse M4 models) further confirm that `Trisicell` inferred trees have phenotypic associations. Moreover, it implies convergent trajectories of melanoma progression. In fact, 10 of the 14 commonly mutated genes in the left subtree of the PDX tree are associated with mitochondria (IDH3B, MRPS35, MRPL9) and protein metabolism (PI4KA, RPSA, SRRM1, WDR18, HSPH1, PSMD7, MDM4), two types of gene function closely linking cell growth with melanocytic differentiation [56].

## 3 Discussion

The large scale of and high levels of noise in single-cell whole transcriptome sequencing (scRNAseq) datasets have limited their use in tumor progression and mutational heterogeneity analysis. Here we introduce `Trisicell`, an algorithmic toolkit to (i) build scalable trees of tumor progression by boosting available tree reconstruction methods to handle large scRNAseq datasets with high levels of noise, (ii) compare multiple trees built for a tumor sample by the use of distinct data types and methods, and establish their consensus tree, and (iii) compute the partition function of a tree of tumor progression to evaluate the robustness of its distinct subtrees and their seeding mutations.

We evaluated `Trisicell` on a tri-modal sequencing dataset comprised of bulk whole exome and transcriptome sequencing data from single-cell derived clonal sublines (each with a homogeneous mutational profile) of a melanoma cell line M4, as well as scRNAseq data from multiple cells from each subline, obtained through Smart-seq2 protocol. `Trisicell` produced robust results on this dataset with a broad consensus among the trees it built for the three data types. Importantly, for the inferred mutation trees, `Trisicell`’s partition function formulation was able to determine that their specific subtrees with distinct growth patterns have very likely been seeded by a distinct set of mutations, suggesting that these mutations are possible subclonal drivers of tumor growth (Figure 4b). In general, the trees strongly correlate with in vivo growth and developmental status, supporting that some of these mutations exhibit function and drive subclonal selection during tumor progression.

We note that only a fraction of the mutation calls on the exome and transcriptome data from the M4 melanoma are shared (see Figure 3d). This may be attributed to the fact that not all genes are expressed, and that post-transcriptional processing and modification can introduce sequence variants not present in the genome [57]. Nevertheless, `Trisicell` inferred trees based on mutation calls using exome and transcriptome data turned out to be nearly identical. This concordance suggests that it is primarily the expressed mutations that are under selection pressure and drive the subclonal expansions in the tumors; other mutations could be neutral or negatively selected under stress. This observation is also supported by the fact that the proportion of mutations that are expressed decreases on lineages from the root to the leaf nodes (see Figure 3b and Figure 3c).

The functionality of expressed mutations are further implied by a comparison between the four major clades of the expressed mutation tree (Figure 3b, c and f) and human melanomas stratified to three classes based on response to anti-CTLA4 therapy by Van Allen et al. [35] (see Figure 5). The significant overlap in the mutated genes between the clades of the mouse bWTS phylogeny and melanoma patients (“sensitive” S1/S2 clades vs. R melanomas and “resistant” S3/S4 clades vs. NR melanomas) provides evidence for convergent (subclonal) expansion across human and mouse melanomas. Noticeably, many of the overlapped genes in the responder sets are similar to the novel tumor suppressor genes (TSG) reported by Cai et al. ([58]), suggesting that they are the bottleneck of subclonal expansion. A comprehensive mutational lineage profiling of human melanomas through the possible use of scRNA sequencing may demonstrate the extent of shared progression histories across tumors with similar treatment outcomes as well as between human and mouse melanomas.

The tri-modal melanoma sequencing dataset we generated for this study is of independent interest, as it offers means to assess the accuracy and robustness of methods for studying the history of tumor progression based on either or both of bulk and single-cell RNAseq data. Additionally, we applied `Trisicell` to analyze six tumors from a preclinical study of M4 melanoma model, among which four were from mice treated with anti-CTLA-4 and the other two from the control treatment. A large set of cells harvested from each tumor were then subjected to Smart-seq2 protocol to produce a large scale scRNAseq dataset. We then applied `Trisicell` to this dataset and compared the tree inferred by the use of mutation calls from single cells to the clusters derived from gene expression profiles. We observed that the subtrees of the tree are better associated with developmental status (as defined by MPS) and neural crest lineage markers than UMAP clusters, which are based on overall gene expression (Figure 6c). Even though the five main subtrees (clades) in the scRNAseq tree all share driver mutations frequently observed in melanoma (i.e. in *Kras*, *Sf3b1*, and *Trp53*), each subtree features a specific combination of mutations and developmental status.

Surprisingly, anti-CTLA-4 treatment did not have an impact on cell lineages. Moreover, among the four anti-CTLA-4-treated tumors, significant subclonal selection occurred in one sample (m122), and adaptation by transcriptional change occurred in the remaining three, consistent with recent studies which suggest that therapeutic resistance is mostly caused by a combination of genetic and non-genetic mechanisms [48]. The comparison we provided between the mouse M4 tree and the mutation tree of a patient-derived xenograft (PDX) from [55] (see Figure 7) provide further evidence of convergent subclonal expansion between the human and mouse melanomas, this time, after anti-CTLA4 therapy. These results demonstrate that `Trisicell` inferred trees allow us to trace lineages and subtrees of cells receiving a particular treatment in detail, adding a key dimension to the studies of intratumor heterogeneity.

In conclusion our results demonstrate the importance of principled, innovative computational methods for studying intratumor heterogeneity and tumor progression. Single cell RNA sequencing has been primarily employed for identifying gene expression differences across distinct cell types. `Trisicell` innovatively uses scRNAseq data to build large scale, robust trees of tumor progression, identify and evaluate distinct subtrees and seeding mutations. As such, it greatly expands possible applications of scRNAseq data to subclonal driver mutation discovery and lineage tracing.

## 5 Methods

### Input data and notation

Given a sequencing experiment involves n single cells (or clonal sublines as in the case of our bWES and bWTS datasets), we denote them by 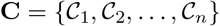. After data processing and mutation calling, suppose that m putative mutations were detected (a mutation is considered to be detected if it is reported as present in at least one single cell) and denote the set of detected mutations by **M** = {*M*_1_, *M*_2_, …, *M*_*m*_}. The observed status of mutations in cells can be represented by a ternary “genotype matrix” *I* with *n* rows (cells), m columns (mutations), where each entry is from the set {0, 1, ?}. Here *I*_*ij*_ = 1 indicates that mutation *M*_*j*_ is reported to be present in cell 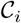 and *I*_*ij*_ = 0 indicates that *M*_*j*_ is reported to be absent in 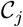. In some cases we do not have sufficient read count information to call any of the present or absent states so we set *I*_*ij*_ =? and call that a “missing entry”. We assume that mutations from regions affected by deletions are filtered from the input and that commonly used infinite sites assumption (ISA) holds, that is, each mutation is acquired (occurs) exactly once and it is never lost in any of the descendants of the cell of its first occurrence.^13^

### 5.1 Trisicell-PartF: a partition function method to assess the likelihood of specific somatic mutations seeding a subclone

We first introduce a partition function formulation for a genotype matrix and describe a novel sampling method to calculate the marginal probability of a user-defined set of cells forming a distinct subtree of the mutation tree of tumor progression, seeded by a user-defined mutation. This formulation allows us to assess our confidence in a particular set of cells to form a subtree/subclone, seeded by a particular mutation.^14^

#### Preliminaries

Let [*x*] = {1, 2, …, *x*} for any positive integer *x*. Recall that we fix *n* and *m* and consider *n* cells, *m* mutations to study matrices *H* ∈ {0, 1} ^*n*×*m*^, where *H*_*i,j*_ = 1 indicates that *i*^th^ cell harbors *j*^th^ mutation. We use *H*_*j*_ to denote the *j*^th^ column of *H*. With a slight abuse of notation, we use binary vectors of length n and subsets of [n] interchangeably. For example *H*_*j*_ denotes the profile of *j*^th^ mutation both as a binary vector and as a subset of cells that harbors *j*. Based on the definition of three-gamets rule from [24], we define 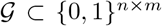 to be the set of matrices that satisfy three-gamets rule. Recall cell lineage trees introduced earlier; we further restrict this definition to *Binary Cell Lineage Tree* (BCLT). BCLTs are cell lineage trees with n observed cells as their leaves and latent subclones as their internal nodes, and each internal node has exactly two children. Notice that any cell lineage tree can be transformed to a BCLT. Also, let BCLT(*H*) denote an arbitrary BCLT consistent with *H* for any 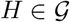. Furthermore, let 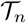 denote the set of all possible BCLTs possible for n cells. We drop the subscript *n* as it is fixed throughout this section. We use bold fonts for random variables to distinguish them from their non-stochastic counterparts. For example we use ****X**** ∈ {0, 1}^*n*×*m*^ to refer to the unobserved ground truth. Finally, we use 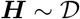 to indicate that the random variable ****H**** follows the distribution implied by some distribution 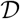.

#### ObtainingP : a matrix of independent probabilities

We define a matrix *P* ∈ [0, 1]^*n*×*m*^ based on the input matrix *I* and noise rates *α* and *β*. The value *P*_*i,j*_ is equal to the probability that *i*^th^ cell harbors *j*^th^ mutation in the ground truth, i.e., ***X***_*i,j*_ = 1, given *I*_*i,j*_, *α* and *β*. Formally, *P*_*i,j*_ = *P*[***X***_*i,j*_ = 1|*I*_*i,j*_, *α*, *β*].

We can use the Bayes rule, along with the assumption that ***X***_*i,j*_ is equally likely to take values 0 or 1 (before observing *I*) to calculate *I* as follows. Fix *α* and *β*, and consider the case *I*_*i,j*_ = 1 first:

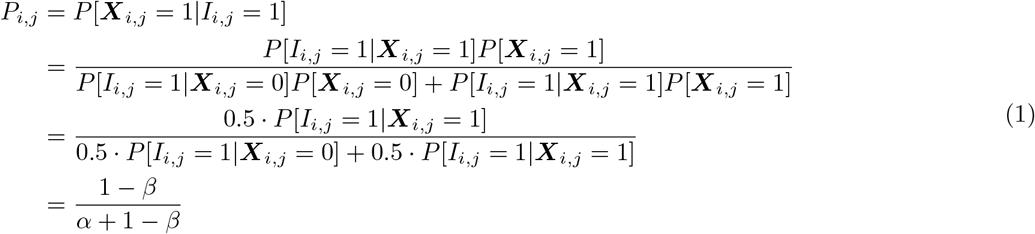

Similarly, for *I_i,j_* = 0 case, we have 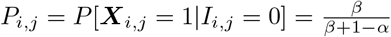.

There are more sophisticated methods for obtaining *P* from the output of SCS process (e.g., see [60]). Our method, presented in this section, is general and works independent of how *P* is calculated.

In the following we present our formulation for partition function problem. Intuitively, the problem aims to measure our confidence for a template of claims about ground truth mutation tree ***X***: Fix a mutation c and a subset of cells R. How confident one can be to claim that the cells in R form a subtree with the root edge of the subtree labeled with c, given P implied by our observations. Notice that this claim is equivalent to ***X***_*c*_ = R.

In the rest of this section we let 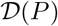 denote the following distribution over {0, 1} ^*n*×*m*^. If 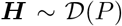 then the probability of ***H***_*i,j*_ = 1 is equal to *P*_*i,j*_, independent of all other coordinates in ***H***.

#### Problem

Given a matrix of independent probabilities *P* ∈ [0, 1]^*n*×*m*^, a mutation *c* ∈ [*m*] and a subset of cells R ⊂ [*n*], estimate 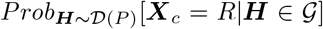. That is, what is the probability that the cth column of ***H*** is equal to *R*, given that ***H*** is coming from the distribution defined by *P* conditioned on being in 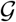.

#### Approach

First, we present a way to sample BCLTs. In short, our algorithm builds a sample tree in a bottom-up manner giving priority to nodes that correspond to the similar rows in *P* as well as nodes that have more mutations in common. Then a number of such samples is constructed and used to estimate 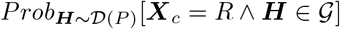 and 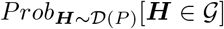, separately. Then our original goal 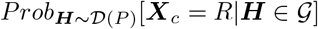 can be calculated via following equation due to the definition of conditional probability:

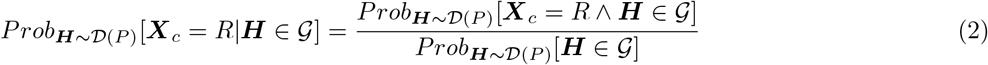

#### 5.1.1 Sampling BCLTs

Algorithm 1 builds a random tree recursively. At each call to this algorithm a pair of rows is chosen based on two criteria: (I) expected number of mutations in common and (II) similarity of the two cells. Then the rows corresponding to this pair of cells are substituted by one new row representing the cells parent. Then the algorithm is recursively called for the new matrix. Let *ɛ* denote the distribution of trees that can be output by this algorithm. Moreover, this algorithm can be modified to output the probability that each tree T is sampled, i.e., Prob_***T***~*ɛ*_ [***T*** = *T*].

**Algorithm 1.**
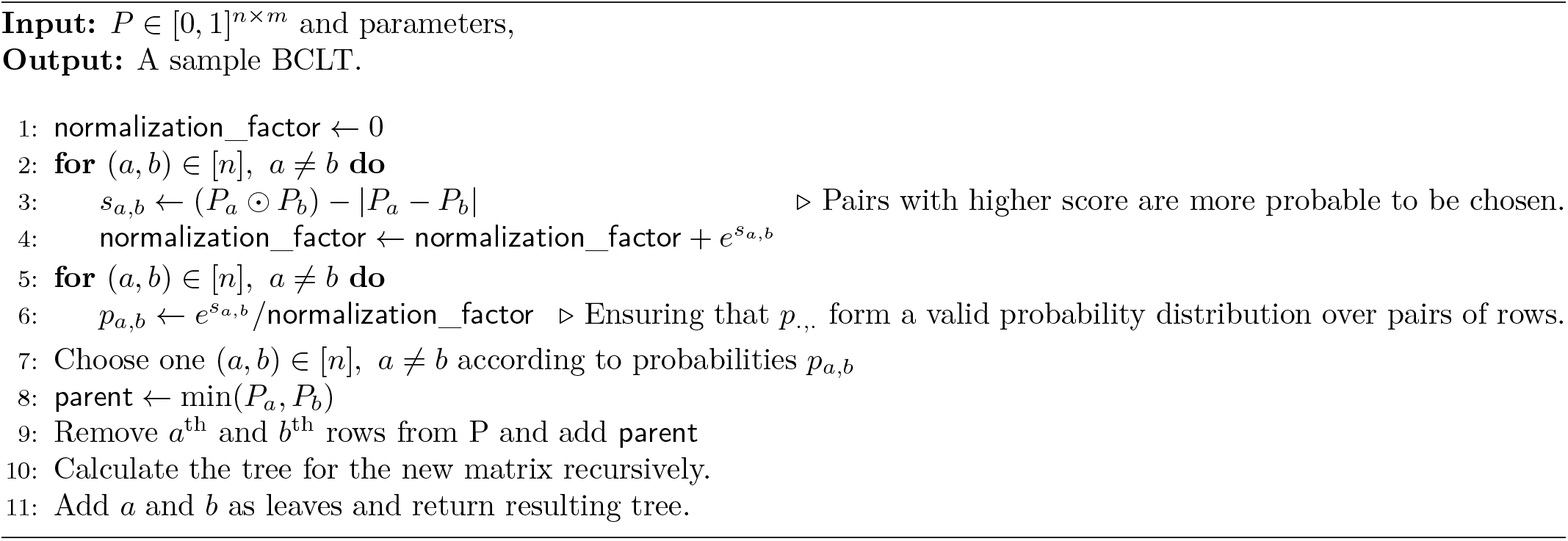
Sample BCLT

#### 5.1.2 Estimating the probability of 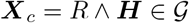

Fix *c, R, A, P*. Given a *T* sampled by Algorithm 1 we define a quantity below that will be used to estimate 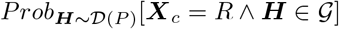:

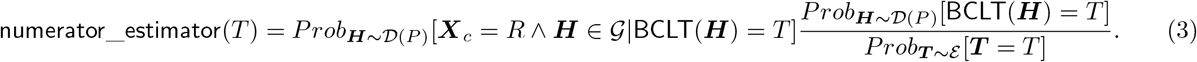

We show in Lemma 5.1 that this value is an unbiased estimator of Equation (2). Algorithms 1 to 3 can output Prob_***T***~*ɛ*_[***T*** = *T*] (along with each *T*), 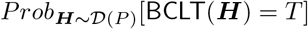, and 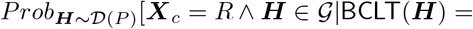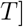, respectively. The proof can be found in Supplementary Section B.

##### Lemma 5.1.

*Let v be the variance of the estimator in Equation (3). Algorithm 4 outputs a value* 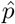 *such that* 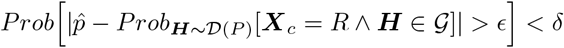*, when N is set to* 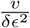.

**Algorithm 2.**
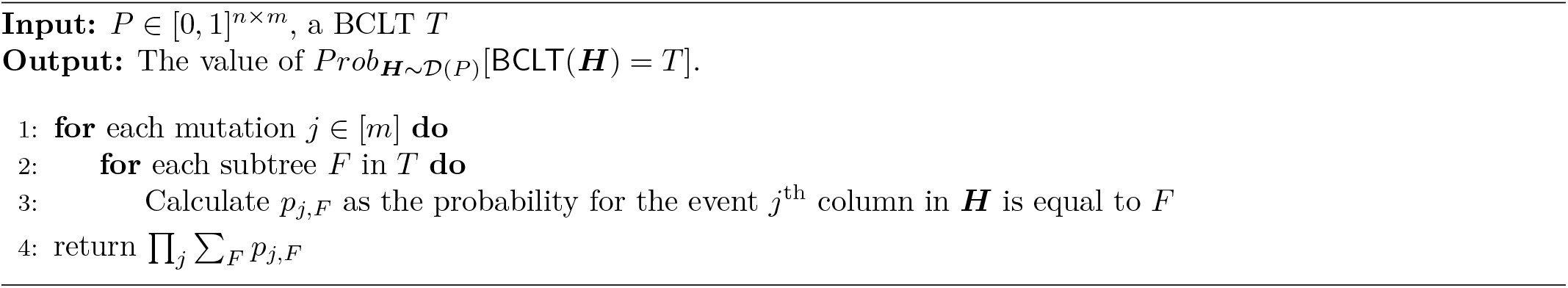
Calculate 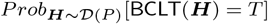 used in Equation (3) given one BCLT

**Algorithm 3.**
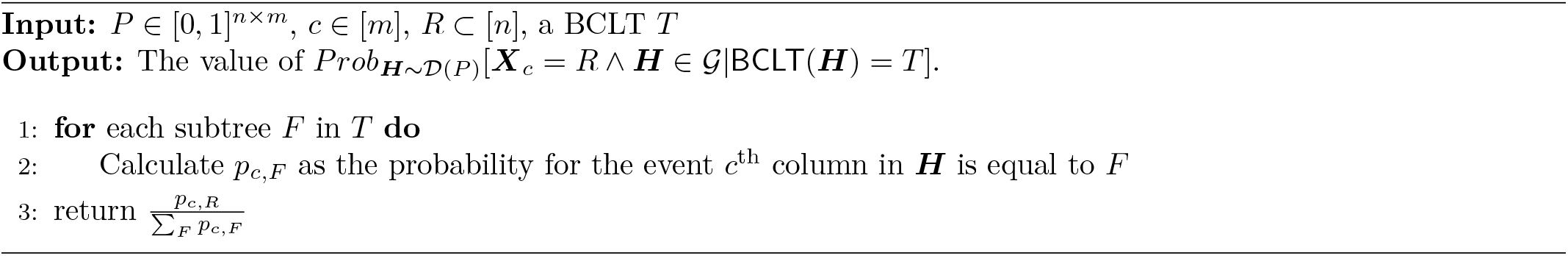
Calculate 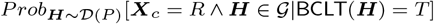 used in Equation (3) given one BCLT

**Algorithm 4.**
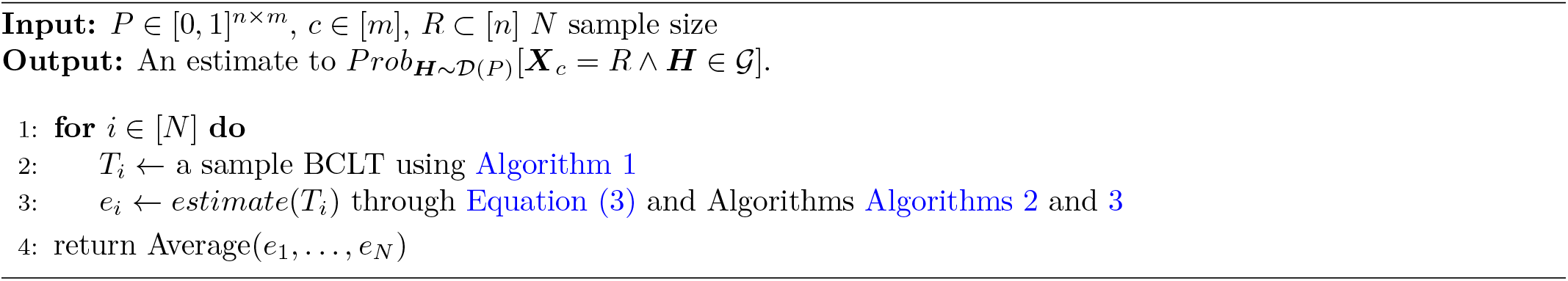
Estimate 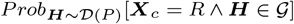

#### 5.1.3 Estimate 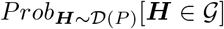

The approach for estimating 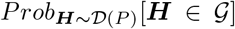 is similar to the previous section, except that we will use the following estimator (instead of numerator_estimator in Equation (3)).

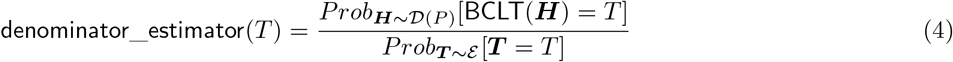

Lemma 5.2 proves a theoretical guarantee for denominator_estimator similar to Lemma 5.1. The proof can be found in Supplementary Section B.

**Algorithm 5.**
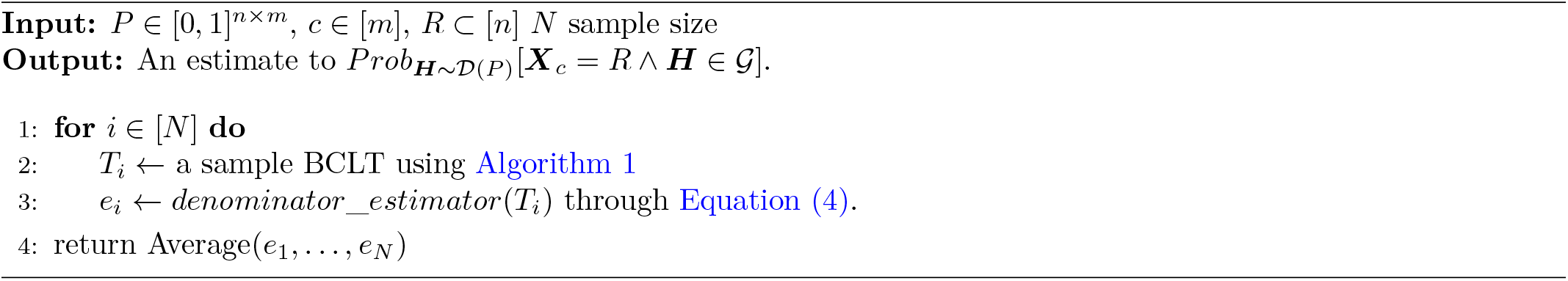
Estimate 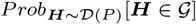

##### Lemma 5.2.

*Let v be the variance of the estimator in* Equation (4). *Algorithm 4 outputs a value* 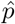 *such that Prob* 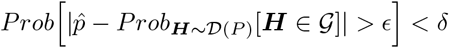, *when N is set to* 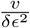.

#### 5.1.4 Putting all together

Given P, c, and R, our algorithm estimates 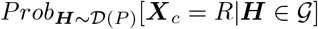 in four phases as follows:

1. Draws a predetermined number of sample BCLTs (Algorithm 1).
2. Estimates 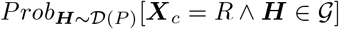 with the guarantee given in Lemma 5.1 (Algorithm 4).
3. Estimates 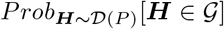 with the guarantee given in Lemma Lemma 5.2 (Algorithm 4).
4. Outputs an estimate for 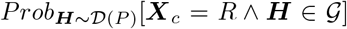 by dividing the estimate from step 2 by the estimate from step 3.

An experimental evaluation of this method on simulated data can be found in Supplementary Section K.

### 5.2 Trisicell-Cons: an algorithm for building consensus of two or more mutation trees

As mentioned earlier, it is desirable to establish a “consensus” between two or more mutation trees of tumor progression. Since the set **C** of cells (or clonal sublines) of the input genotype matrix representing a tumor is fixed but the mutation calls may vary, we focus on establishing a consensus between cell lineage trees rather than mutation trees.^15^ Thus we introduce a simple, deterministic, exact algorithm for building a consensus tree of two or more cell lineage trees, which runs in time quadratic with the input size. The ancestor-descendant relationships between cells implied by the consensus of multiple cell lineage trees, each obtained through distinct methods and data types, are likely to be more reliable. Edges maintained in the consensus tree are also more likely to be labeled by mutations that “seed” the associated subtree.

Given sequencing data from n single cells, labeled 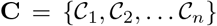 *T*^*a*^ and *T*^*b*^ denote two rooted trees depicting lineage relationships between the sequenced cells. For each of the two trees, we assume that: (i) each edge has a positive “weight”, (ii) each node has 0, 1 or more labels and each leaf node has at least 1 label, (iii) each internal node without a label has at least two children, (iv) each label occurs exactly once in each tree, (v) the set of labels are identical in the two trees.^16^ The weight of an edge represents the number of mutations that distinguish a (possibly inferred) parent node from its child. We call such a tree a *generalized cell lineage tree* and as the label sets of the two trees are identical, we say they are *comparable*.

We now define the *merge-with-parent* operation on any given non-root node *v* of a tree: the operation merges *v* with its parent *w* and deletes edge (*w*, *v*) so that (a) the new label set of w becomes the union of the original label sets of *w* and *v*, and (b) the new set of children of *w* is the union of the original sets of children of *w* and *v* minus *v* (as it ceases to exist). Observe that after *merge-with-parent* operations, the sets of labels on each node remain disjoint from one another. The *cost* of a merge-with-parent operation is defined to be the weight of the edge (*w*, *v*). Our goal is to perform a sequence of merge-with-parent operations on the two trees with minimum total cost so that the resulting trees are *isomorphic* – and can be thought of as a single tree *T*^*c*^, which we call *the consensus tree* of *T*^*a*^ and *T*^*b*^.

#### Definition 5.1.

*Given a pair of comparable cell lineage trees and a node v in either tree, let L(v) denote the complete set of labels in the subtree rooted at v (including the labels of v itself). Then the two trees are isomorphic, if there is a bijection between their nodes such that for any pair of matched nodes v and w, L(v) = L(w)*.

For any two nodes *v*, *w* in one tree, *L*(*w*) ⊆ *L*(*v*) if w is a descendant of *v*, and, if there is no ancestor/descendant relationship between *v* and *w*, *L*(*v*) and *L*(*w*) are disjoint. Also, note that a matched leaf in *T*^*a*^ must be matched to a leaf in *T*^*b*^, and their labels must be identical. This isomorphism implies that the two trees look identical:

#### Lemma 5.3.

*Any two comparable cell lineage trees T ^a^ and T ^b^ are isomorphic if and only if there is a bijection between their nodes where (i) the label sets of matched nodes are identical, and (ii) children of matched nodes are also matched.*

The proofs of Lemma 5.3 and the following lemmata in this section can be found in Supplementary Section C. We now describe our algorithm to compute *T*^*c*^. For simplicity, we consider a node to be its own lowest ancestor.

#### First Step

For any pair of labels *x* and *y* where *x* is in an ancestor of the node of *y* in *T*^*a*^, but not in *T*^*b*^, the algorithm performs subsequent merge-with-parent operations in *T*^*b*^, from the node of *x*, up to the lowest common ancestor of the nodes of *x* and *y* so *x* becomes an ancestor of *y* in *T* ^*b*^. (A symmetric action happens when *T*^*a*^ and *T*^*b*^ are reversed.) Note that, if *x* is a proper ancestor of *y* (i.e., they are not in the same node) in *T*^*a*^ and *y* is a proper ancestor of *x* in *T*^*b*^, the sequences of merge operations triggered in the two trees will end up placing *x* and *y* in the same node in both *T*^*a*^ and *T*^*b*^. We now show that the above operations are necessary.

##### Lemma 5.4.

*Let *x* reside in an ancestor of the node containing *y* in *T*^a^ (for simplicity, we can say *x* is an ancestor of *y*) and not in *T*^*b*^. Let *z* be the lowest common ancestor or *x* and *y* in *T*^b^. Then, any construction of *T*^c^ must involve removing the edges between *x* and *z* in *T*^b^ through merge-with-parent operations.*

We can see that once the first step concludes, ancestor-descendant relationships are consistent across the two trees.

##### Lemma 5.5.

*After the first step, any two labels *x* and *y* (a) are either in the same node in both trees, or (b) have the same ancestor-descendant relationship in both trees, or (c) have no ancestor-descendant relationship in either tree.*

The output of the first step will thus be two comparable cell lineage trees in which no pair of labels have “different” ancestor-descendant relationship between the trees. We will call them *T*^′*a*^ and *T*^′*b*^.

#### Second Step

Based on the observation above, the algorithm now replaces each node’s label set with a single new label. The result of this step will be two trees *T*″^*a*^ and *T*″^*b*^, respectively corresponding to trees *T*″^*a*^ and *T*′^*b*^, in which (1) each leaf node has exactly 1 label and each internal node has either 1 or 0 labels, where each label represents a set of labels from the original trees; (2) the label set of the two trees are identical. As such, *T*″^*a*^ and *T*″^*b*^ are (still) comparable cell lineage trees that satisfy the following.

##### Lemma 5.6.

*For each leaf v in a tree, there exists exactly one leaf w in the other tree where L(v) = L(w). For each internal node v in each tree there exists at most one internal node w in the other tree such that L(v) = L(w)*.

We call a node *v* in each tree for which there is a corresponding node *w* in the other tree such that *L*(*v*) = *L*(*w*) a *matched* node. All other nodes are said to be *unmatched*. The following holds by definition.

##### Lemma 5.7.

*If all nodes of any of the trees are matched and the trees have the same number of nodes then the two trees are isomorphic.*

##### Lemma 5.8.

*For each tree, any node v with a label l must have a matched node w in the other tree with label l*.

#### Third Step

The algorithm processes *T*″^*a*^ and *T*″^*b*^ to compute their consensus *T*″^*c*^, which is convertible to *T*^*c*^. W.l.o.g., consider any matched internal node *v* in *T*″^*a*^ for which the subtree rooted at *v* has no matched internal nodes (including when all children of *v* are leaves). Let *w* be the node matched to *v* in *T*″^*b*^. The algorithm computes the consensus of the subtrees rooted at *v* and *w* (these subtrees must be comparable cell lineage trees) by performing merge-with-parent operations on all internal nodes such that the subtrees end up as *star trees*; a star tree is a rooted tree with no internal nodes.

In case *v* and *w* are the roots of respective trees *T*^*a*^ and *T*^*b*^ the algorithm terminates at this point since the resulting trees must be isomorphic.^17^ If they are not roots, then the algorithm “replaces” the subtrees rooted by *v* and *w* by respective leaf nodes *v* and *w*, both with a new label *l* and iterates.

##### Lemma 5.9.

*The algorithm described above computes a consensus tree T ^c^ in ((|T ^a^| + |T ^b^|)*^2^) *time and linear space.*

##### Lemma 5.10.

*The algorithm terminates with isomorphic trees.*

We can now argue about the optimality of the algorithm.

##### Lemma 5.11.

*Each merge-with-parent operation by the algorithm is necessary for obtaining a consensus tree.*

Since all the edges that the algorithm removes are those that must be removed, the Corollary below follows.

##### Corollary 5.1.

*The algorithm terminates with a consensus by removing edges whose total cost is minimum possible.*

##### Corollary 5.2.

*The algorithm can be generalized to k* > 2 *input trees by recursively constructing the consensus tree of the first k* − 1 *input trees and then building the consensus of that tree and the final input tree.*

### 5.3 Trisicell-Boost: a divide and conquer booster for scalable inference of the history of tumor progression

Due to false mutation calls and missing entries, the observed matrix *I* is usually different from the true genotype matrix and, as such, it does not perfectly match any mutation tree of tumor progression. In other words, there does not exist a mutation tree such that, for each single cell its observed genotype matches genotype of some of the tree nodes (see [20] for more details). There exist several methods that aim to *correct* (and possibly impute missing) entries of *I* so that the resulting corrected matrix *Y* perfectly matches some mutation tree, which is then reported as a tree of tumor progression of the sequenced tumor. When searching for *Y*, the existing methods aim to minimize the weighted number of 1 → 0 “flips” (which represent false positives) and 0 → 1 “flips” (which represent false negatives) required to convert *I* to *Y* [7, 17, 23, 24, 25].

Let 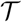 denote (one of) the most likely mutation tree(s) of tumor progression, where the log-likelihood of a tree is defined based on a scoring model commonly used in the existing tree inference methods. While the existing methods such as PhISCS or SCITE can in principle infer 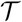, these methods do not scale well in terms of the running time on large datasets. As we show, for example, on simulated genotype matrices of size 1000 × 1000, these tools either do not terminate within a time limit of 24 hours (e.g. PhISCS, ScisTree) or yield trees that have a sizeable number of mutational pairs that are misplaced in terms of commonly used ancestor-descendant and different-lineage measures (e.g. SCITE). This motivated us to introduce `Trisicell-Boost`, a novel method for reconstructing mutation trees of tumor progression that can handle such (simulated) datasets within the time limit and achieve high accuracy. As will be shown, `Trisicell-Boost` is robust to various types of noise, including the presence of mutations affected by copy number aberrations, possibly resulting in ISA violations. On “real” melanoma WES data involving clonal sublines, the `Trisicell-Boost` inferred trees are not only (almost) identical to that obtained by (much slower but provably optimal) PhISCS-BnB, but also very similar to that it obtains on matching WTS data. Finally, these trees, as well as the tree obtained on a melanoma scRNAseq dataset, highly associate with phenotypes such as cell differentiation status and tumor growth.

In designing `Trisicell-Boost` we extensively rely on the observation that any subset of mutations from **M** also forms a mutation tree [61]. We first sample a given number of submatrices of I, each obtained by taking a random subset of z columns (mutations) of I. In our experiments we sample 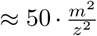 submatrices, implying that, on average, each pair of mutations appears together in approximately 50 submatrices. This plays an important role in obtaining reliable *weights* that are defined and used below. Although in principle the constant 50 can be any other positive value depending on computational resources and time, we recommend using at least 10. Similarly, we recommend that each sampled submatrix has at least 10 columns (i.e., z ≥ 10). We then run PhISCS on each of the submatrices and obtain a mutation tree for the corresponding set of sampled mutations. Note that on the smaller input matrices, PhISCS, under the model that it uses, can provide a guarantee of optimality of the reported solutions. This is the main reason why it was our primary choice for obtaining trees on sampled submatrices, although any other tool that performs well on smaller inputs (e.g., SCITE, OncoNEM, ScisTree or several others) can be used for this purpose.

Once the tree for each sampled submatrix is obtained, we do the following: for each pair of mutations *M*_*i*_ and *M*_*j*_ and each submatrix in which both *M*_*i*_ and *M*_*j*_ were sampled, we obtain *(tree) dependency* between them. Given a submatrix where both are present, *M*_*i*_ and *M*_*j*_ can have one of the following dependencies in the inferred tree:

1. **Ancestor-descendant** where (the first occurence) node of *M*_*i*_ is an ancestor of that of *M*_*j*_. We denote by a_*ij*_ the total number of submatrices for which *M*_*i*_ and *M*_*j*_ have this dependency in their inferred trees.
2. **Descendant-ancestor** where node of *M*_*i*_ is a strict descendant of the node of *M*_*j*_. As per above, we denote the number of submatrices for which this is a case as *d*_*ij*_. Note that *d*_*ij*_ = *a*_*ji*_.
3. **Same-node** where both *M*_*i*_ and *M*_*j*_ occur (for the first time) at the same node. We denote the number of reported trees in which this holds by *s*_*ij*_.
4. **Different-lineage** where *M*_*i*_ and *M*_*j*_ occur at nodes that are on different tree lineages (i.e., their nodes are different and neither of them is an ancestor of the other). We denote by l_*ij*_ the number of submatrices for which *M*_*i*_ and *M*_*j*_ are in this dependency in their inferred trees.
5. **Unknown dependency** where the tree inference tool used reports that at least one of *M*_*i*_ and *M*_*j*_ is absent in all cells.

Let *t*_*ij*_ = *a*_*ij*_ + *d*_*ij*_ + *l*_*ij*_ + *s*_*ij*_. Based on the above, we define three *weights*: *A*_*ij*_ (*ancestor-descendant weight*), *d*_*ij*_ (*descendant-ancestor weight*), and *L*_*ij*_ (*different-lineage weight*), which are respectfully defined as:

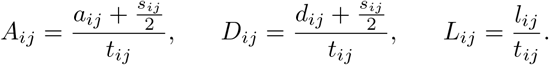

Consider now an arbitrary mutation tree T of size m + 1 where the root node is mutation-free representing a population of healthy cells, at each node we have exactly one mutation label from **M** and no mutational label appears more than once. We say that mutation *M*_*i*_ is an ancestor of *M*_*j*_ if and only if node labeled by *M*_*i*_ is an ancestor of node labeled by *M*_*j*_. Analogously we define *M*_*i*_ being descendant of or on different lineages compared of *M*_*j*_. Using the above notation we can define score *S*(*T*) of the tree *T* as follows:

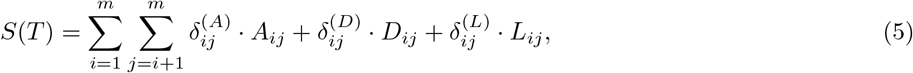

where 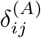 is an indicator variable with value set to 1 if and only if mutation *M*_*i*_ is ancestor of mutation *M*_*j*_ in *T*. Variables 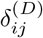 and 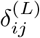 are defined analogously and indicate whether *M*_*i*_ is descendant of *M*_*j*_ (variable 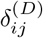) or whether these two mutations occur on different lineages (variable 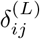).

We leave finding a tree *T* for which *S*(*T*) is maximized as an open problem and instead propose a fast heuristic for building a tree iteratively by adding mutations in a specified order and one at a time. Full details of this heuristic are provided in Supplementary Section D.

### 5.4 Wet lab data generation to validate Trisicell

The “M4” mouse model of UV-induced melanoma used in this study is described in [32]. In brief, pups of Hgf-transgenic C57BL/6 mice received UV irradiation at post-natal day three, and melanoma occurred in 8-12 months in some mice. One of the tumors was harvested from a mouse and subjected to in vitro culture. After consecutive passage, the melanoma line was built and named B2905 cell line.

#### 5.4.1 Establishing and characterization of the 24 clonal sublines of B2905

The parental B2905 cells were subjected to fluorescence-aided cell sorting (FACS) to sort single cells into individual wells of a 96-well plate, which was brought to in vitro culture. Cells were found growing in twenty-four wells. They were expanded by consecutive passages to build the twenty-four clonal sublines derived from single cells of B2905. To characterize each subline, exome and single cell RNA were analyzed. For exome analysis, genomic DNA was prepared from cells harvested from each subline and subjected to exome capture protocol, followed by whole exome sequencing on Illumina HighSeq platform. For single cell analysis, 16 single cells from each subline were sorted to individual well of 96-well plate by FACS. cDNA library was prepared from the RNA of single cells in each well following Smart-seq2 protocol [33]. After quality control check, the cDNA libraries of 8 selected cells from each subline (in total 8×24 = 192) were sequenced on the NovaSeq platform. For downstream analysis and mutation calling, 175 cells that had sufficient coverage across all genes were selected.

#### 5.4.2 Analysis of in vivo tumors derived from the clonal sublines of B2905

To test the growth kinetics in vivo, one million cells of each clonal subline were implanted into five syngeneic C57BL/6 mice subcutaneously. The size of tumors was measured twice a week. Eleven sublines showed 100% tumor-take rate within 90 days. Three tumors from each of these sublines were selected as triplicates. They were subjected to RNA preparation, followed by mRNA sequencing.

#### 5.4.3 single cell sequencing of melanoma in the preclinical study

For the preclinical study of anti-CTLA-4, one million cells of B2905 were subcutaneously implanted into twenty C57BL/6 mice. When the tumors reaching 75 mm^3^, the mice were randomized to two groups of equal number to receive either mouse anti-mouse CTLA-4 antibody (clone 9D9, Bio X Cell) or mouse IgG2b as isotype control (clone MPC-11, Bio X Cell). When the tumor reached around 500 mm^3^, the mouse was euthanized for tumor harvest. The two control group tumors were harvested at day 41 and 54 and from anti-CTLA-4 group, one tumor was harvested at day 54, another at day 75 and final two at day 85 (see Supplementary Section G for tumor size detail). A single cell suspension was prepared from the tumor as described previously [62], stained with antibodies, nuclear dye, and viability dye, and then subjected FACS to sort CD45^−^;CD31^−^ nucleated alive single cells into individual wells of 96-well plates. A total of 1008 cells from four anti-CTLA-4 treated tumors and 384 cells from the control tumors were collected. cDNA library was prepared from the RNA of single cells in each well following Smart-seq2 protocol [33]. After a quality control check, the cDNA libraries from 1248 cells (960 from four anti-CTLA-4 treated tumors and 288 cells from the control tumors) were sequenced on the NovaSeq platform. For downstream analysis and mutation calling, 764 cells (595 from four anti-CTLA-4 treated tumors and 169 cells from the control tumors) that had sufficient coverage across all genes were selected.

### 5.5 Somatic variant calling pipeline

Several studies have explored SNVs/Indels in scRNAseq data to decipher the heterogeneity of cell populations [63, 64, 65, 66, 67]. We used GATK Best Practices [68] to call mutations from scRNAseq, bWTS and bWES datasets.^18^ Briefly, after trimming and quality control of reads by FastQ (v0.11.9) [69] and Trim Galore (v0.6.5), raw reads were mapped to the mouse genome (GRCm38, mm10) using STAR (v2.7.3a) [70] in two-pass mode which improves the mapping quality at junction sites. Then Picard (v2.22.3) was used for removing duplicated mapped reads and doing indel realignment and base re-calibration. During base quality recalibration, dbSNP variants were used as known sites. In the next step, HaplotypeCaller (v3.8) [71] was used to call the mutations for every single cell separately in GVCF mode. After that, GenotypeGVCFs was used to call the mutations in a joint mode (as pseudo-bulk). These steps keep specificity and sensitivity of calling mutations high. Afterwards, the mutations were annotated using ANNOVAR (v2019.10.24) [72]. Following that, mutations that marked as unknown and non-exonic were filtered out. For further confidence, mouse SNPs/Indels filtration was applied to the VCF files based on normal mouse spleen SNPs in addition to a curated in-house database. Finally, somatic-mutations associated with less than 10 reads in at least 2 cells were filtered out. (see Supplementary Section N for more details)

### 5.6 single cell gene expression quantification and quality control

RSEM (v1.3.2) [73] was used to extract expected counts, TPM and FPKM information. The Seurat package (v4.0.0) [49] was used for clustering and UMAP analysis. The low-quality cells were removed if any of the following is true: 1) percentage of mitochondrial genes is above 20%; 2) the number of reads is less than 500,000; 3) nFeature_RNA < 1000 (see Supplementary Sections G and M for more details)

To remove batch effect due to different datasets, we used the functions of `FindIntegrationAnchors` and `IntegrateData` in Seurat package. Clustering analysis was performed with `FindClusters` (resolution = 0.5).

### 5.7 Differential gene expression, pathway, and gene set enrichment analyses

Differential gene expression was performed using DESeq2 (v1.30.0) [74]. The contrasts include aCTLA4 cells vs IgG cells in each branch in the mutation tree. FDR was used as 0.1.

The rank list of differentially expressed genes (DEGs) was used to perform gene set enrichment analysis (GSEA) with the R package FGSEA (v1.16.0) [75]. We performed GSEA with the Hallmark gene sets available in the R package MSigDB (v7.2.1) [76]. Results are presented in Supplementary Section F.

## Data availability

Sequence data have been deposited at NCBI GEO with the accession code GSExxxxxx.

## Code availability

Trisicell is available at https://trisicell.readthedocs.io

## Acknowledgements

We thank Maria Hernandez, Kimia Dadkhah and Yongmei Zhao (at Frederick National Laboratory for Cancer Research) for helping us sequence our samples. This work is supported in part by the Intramural Research Program of the National Institutes of Health, National Cancer Institute and utilized the computational resources of the NIH Biowulf high performance computing cluster (http://hpc.nih.gov) and Gurobi (http://www.gurobi.com) to solve some optimization problems. F.R.M. was supported in part by Indiana U. Grand Challenges Precision Health Initiative.

## Footnotes

1 Note that our focus in this work is on mutation trees (see [5] for formal definition). In all figures provided in this work, we provide compacted mutation trees obtained by merging together mutations that have same inferred presence-absence status in all single cells (as relative placement of such mutations is indistinguishable from the data).

2 Consider a simple example with a noise-free input comprised of three cells and *m >* 4 mutations. Suppose the first cell has only the first mutation, the second cell has only the second mutation and the third cell has all mutations except the first mutation. Among these cells, the “nearest” two are the first and second, with a Hamming distance of 2. A distance-based greedy method could conclude that the first and second cells are siblings, and their parent is a sibling of the third cell. ISA dictates that this parent node does not include any mutation. This would imply that the second mutation occurs independently both in the third cell and the second cell, violating ISA. However there is an alternative, ISA satisfying solution where the second and third cells are siblings, and their parent is a sibling of the first cell.

3 In the context of RNA-structure prediction, given the collection of potential base pairs as a set of random variables, the partition function represents the marginal probability of a particular base pair (or set of “non-conflicting” subset of base pairs), i.e., the total probability of all possible configurations that feature the base pair(s).

4 Examples include the number of ancestor-descendant and different-lineage relationships shared across the trees compared, as well as more sophisticated measures such as the Multi-Labeled Tree Edit Distance and others [24, 39, 40, 41, 42].

5 which is important especially since many methods build trees based on mutations shared by at least two cells or pre-cluster cells based on their mutational profiles

6 Another parameter used in our simulations is the missing value rate *γ*, which determines the fraction of entries in the genotype matrix which are unknown. We used *γ* = 0.2 in our simulations. This choice is consistent with the missing value rates observed in single-cell genome sequencing data but is somewhat lower than that we observed in our real scRNAseq data. Our simulations aim to compare the performance of `Trisicell-Boost` to the methods it boosts, in particular SCITE and PhISCS, which were developed for genome sequence data. For higher missing value rates, a comparison of `Trisicell-Boost` to SCITE, PhISCS, as well as DENDRO and Cardelino, tools developed specifically for RNAseq data, is provided for real scRNAseq datasets.

7 Since the model employed by SCITE and PhISCS are identical, the trees they built were isomorphic for many of the smaller datasets; thus we do not report `Trisicell-Boost`’s results for SCITE and PhISCS separately.

8 Even though for most sublines, the expression profiles of all three replicates were very similar, in rare cases a replicate had a substantially different gene expression profile, suggesting an epigenomic alteration.

9 Survival time is defined as the post-inoculation time required for the tumor to reach the pre-determined endpoint, size of 1000 mm^3^.

10 bWTS data does not include all mutations or sublines.

11 The tumors were harvested when their size reached about 500 mm^3^ as endpoint. For scRNAseq analysis, the two control group tumors were harvested at day 41 and 54. From anti-CTLA-4 group, one tumor was harvested at day 54, another at day 75 and final two at day 85 (see Supplementary Section G). The variable time of endpoint demonstrates the diverse growth kinetics and therapeutic responses of this model.

12 Briefly, Seurat uses graph-based clustering approaches such as [50, 51] that embed cells in a graph structure where edges between cells indicate similar feature expression patterns. It then attempts to partition this graph into interconnected “communities” using Louvain or smart local moving (SLM) methods [52].

13 Nevertheless, we have assessed the robustness of our methods in the presence of undetected deletions that result in mutational losses and possible ISA violations.

14 The formulation naturally generalizes to any user defined set of mutations - including the empty set, in which case it represents the probability of a user defined set of cells to form a subclone without a specific seeding mutation.

15 In addition, the problem of building a consensus across mutation trees through “edit operations” including deletion or move of mutation labels from one edge to another is NP-hard [24].The single polynomial time computable “consensus” construction approach allows edit operations on leaf nodes only [39].

16 The trees could have been built by two distinct methods on the same set of mutation calls on each single cell, or could have been built by the same method using different subsets of mutations.

17 Note that at the time of termination, all subtrees that have been replaced by a single leaf node should be reinstated.

18 Depends on the data there are some modifications in the pipeline.

